# Protein interactions, calcium, phosphorylation, and cholesterol modulate CFTR cluster formation on membranes

**DOI:** 10.1101/2024.05.03.592454

**Authors:** Yimei Wan, Rhea Hudson, Jordyn Smith, Julie D. Forman-Kay, Jonathon A. Ditlev

**Affiliations:** Department of Biochemistry, University of Toronto, Toronto, Ontario, Canada; Program in Molecular Medicine, Hospital for Sick Children, Toronto, Canada; Program in Cell Biology, Hospital for Sick Children, Toronto, Canada

## Abstract

The Cystic Fibrosis Transmembrane Conductance Regulator (CFTR) is a chloride channel whose dysfunction leads to intracellular accumulation of chloride ions, dehydration of cell surfaces, and subsequent damage to airway and ductal organs. Beyond its function as a chloride channel, interactions between CFTR, ENaC, and SLC transporter family membrane proteins and cytoplasmic proteins, including calmodulin and NHERF-1, co-regulate ion homeostasis. CFTR has also been observed to form mesoscale membrane clusters. However, the biophysical mechanisms that regulate the formation of CFTR clusters are unknown. Using a combination of computational modeling and complex biochemical reconstitution assays, we demonstrate that multivalent protein-protein interactions with CFTR binding partners, calcium, and membrane cholesterol can induce CFTR cluster formation on model membranes. Phosphorylation of the intracellular domains of CFTR also promotes cluster formation in the absence of calcium, indicating that multiple mechanisms can regulate CFTR cluster formation. Our findings reveal that coupling of multivalent protein and lipid interactions promote CFTR cluster formation consistent with membrane-associated biological phase separation.

**SIGNIFICANCE STATEMENT:** Mutations in the cystic fibrosis transmembrane conductance regulator (CFTR) membrane protein underlie cystic fibrosis. It is thought that molecular “hubs” of CFTR and its binding partners co-regulate ion homeostasis and that disruption of these clusters can result in disease. However, the physical basis for molecular hub formation is unclear. In this study, we present evidence that multivalent protein and lipid interactions drive the formation of mesoscale CFTR-containing clusters or “hubs” on model membranes in a manner consistent with biological phase separation. These data provide important insights into physical mechanisms that modulate CFTR membrane organization and offer a new lens for the development of corrective therapeutics.

## INTRODUCTION

The proper organization of receptors and channels on plasma membranes is essential for cellular signaling pathway regulation and ion homeostasis [1,2]. Biomolecular condensates are widely recognized for their ability to functionally organize the nucleoplasm, cytoplasm, and membranes [3–7]. Phase separation driven by multivalent interactions between proteins and nucleic acids can facilitate the formation of condensates [8,9]. Posttranslational modifications, such as phosphorylation or glycosylation, are known to alter their formation, properties, and functions [10–12]. Lipids in the membrane can also undergo phase separation into cholesterol-rich or -depleted domains [13,14]. Many membrane-associated proteins that can undergo phase separation are localized to cholesterol-rich regions [15,16], suggesting a link between protein and lipid phase separation. Indeed, recent studies involving the T cell signaling protein, LAT, and the B cell receptor demonstrated the importance of coupled lipid and protein phase separation for the formation of functional protein clusters on membranes [17,18].

The Cystic Fibrosis Transmembrane Conductance Regulator (CFTR) is a chloride channel of the ATP-binding cassette (ABC) transporter superfamily that enables the movement of chloride ions down their electrochemical gradient across the plasma membrane [19–23]. CFTR channel opening is regulated by ATP binding to the intracellular nucleotide binding domains 1 and 2, and by phosphorylation of the ∼200-residue regulatory (R) region, following activation of the cAMP pathway [24–30]. The R region is an intrinsically disordered segment of the cytoplasmic portion of CFTR that, along with the intrinsically disordered C-terminal region, mediates numerous intra- and intermolecular interactions that regulate CFTR localization on the plasma membrane and its function [31–34].

Beyond its function in chloride homeostasis, CFTR is a molecular “hub” that modulates cellular ion homeostasis by co-localizing other ion transporters and channels, such as SLC26 family transporters and epithelium sodium channel (ENaC), and intracellular signaling proteins, such as NHERF-1 and calmodulin, through multivalent interactions [32,35–37] (Figure 1). Within these “hubs,” CFTR and its binding partners co-regulate one another. However, clearly defining the mechanism(s) of co-regulation between CFTR and its binding partners is often challenging due to their observed complexity. For instance, CFTR is inhibitory of ENaC in most cells [36,38–41] but can activate ENaC in sweat ducts [42,43]. ENaC is generally accepted to activate CFTR [44], although one study suggests an inhibitory role [45]. For SLC26A3 and SLC26A6, the R region–STAS interaction is stimulatory for both CFTR and SLC26 transporter activity [35]. Alternatively, SLC26A9 activates CFTR [46], while CFTR can either activate or inhibit SLC26A9 [47]. These examples demonstrate that the co-regulation between CFTR and its partners is a complex process; studies on co-regulation are often contradictory and little is understood about the underlying molecular mechanisms that contribute to co-regulation. Nevertheless, it appears that protein activity is intimately tied to the co-localization in protein-dense clusters on membranes, sometimes referred to as “macromolecular” clusters. Indeed, CFTR and its binding partners can form micron scale clusters on plasma membranes [48–50]. Treatment of cells with inhibitors of calcium uptake by the endoplasmic reticulum, thapsigargin and curcumin, which increases calcium concentration in the cytoplasm, resulted in increased CFTR cluster size and number on the plasma membrane [50–52]. Calcium also activates PKA and PKC, which can phosphorylate the CFTR R region and modulate its association with other proteins [31,32]. Other research suggests a role for membrane cholesterol in CFTR cluster formation; depletion of cholesterol reduced cluster formation while increased cholesterol density resulted in increased CFTR cluster size and number [49]. While CFTR clustering is known to be modulated by calcium levels and the density of membrane cholesterol, the physical mechanisms that drive CFTR clustering and the role of specific protein interactions and phosphorylation are unclear.

**Figure 1.**
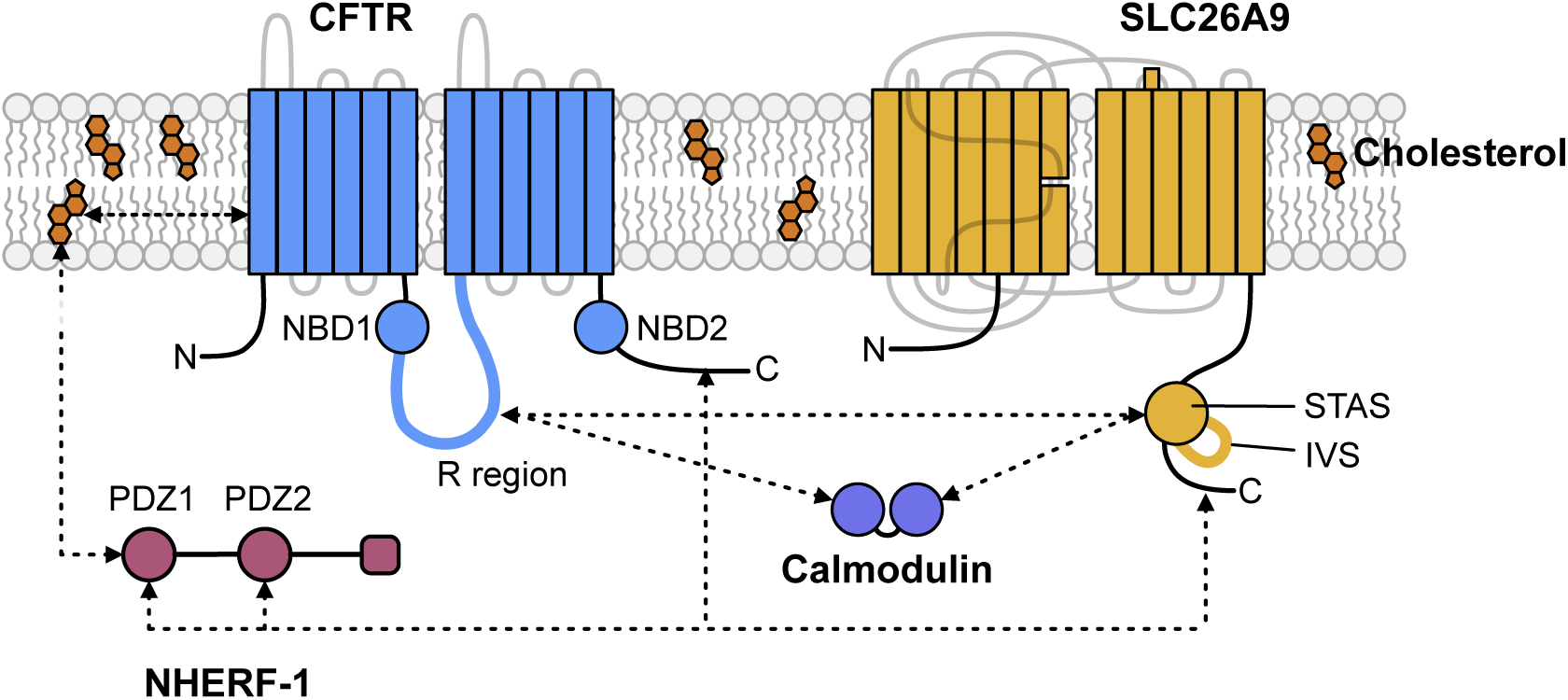
CFTR is a multivalent hub for macromolecular complexes on membranes. The CFTR transmembrane domains interact with cholesterol in the membrane, CFTR R region contains multiple calmodulin and SLC26A9 binding sites, and the C terminal tail of CFTR binds NHERF-1. Calmodulin can interact with CFTR and SLC26A9. NHERF-1 can interact with membrane cholesterol, SLC26A9, and CFTR. SLC26A9 can interact with CFTR, NHERF-1, and calmodulin.

Mutations to CFTR directly cause Cystic Fibrosis (CF) by reducing the amount of CFTR at the plasma membrane, improper gating of the channel, and/or inefficient ion transport by CFTR [53,54]. In addition, the most common mutation to CFTR, the deletion of phenylalanine at position 508 (F508del) results in defects in cluster formation on the plasma membrane [50]. A previous study found that the F508del mutation can prevent dimerization of the first CFTR nucleotide binding domain, thus preventing a self-associating interaction that could contribute to cluster formation [55]. However, it is unclear if the lack of CFTR cluster formation is due to mutations abrogating interactions that may contribute to cluster formation, the density of CFTR successfully trafficked to the cell surface being too low for cluster formation, or a combination of both.

Membrane-associated phase separation underlies the formation of numerous protein-rich clusters that can depend on posttranslational modifications and organization of membrane lipids, including cholesterol [1,16]. Because CFTR “macromolecular” clusters are composed of multivalent proteins, we reasoned that phase separation may underlie cluster formation. This possibility is especially intriguing because phase separation enables a physical understanding of CFTR and binding partner cluster formation and provides a lens through which the aforementioned complexity of CFTR co-regulation in cells can be appreciated [3]. However, it is unclear whether multivalency and lipid domain formation can promote CFTR cluster formation on membranes. In this study we use *in silico* computational modeling and *in vitro* biochemical reconstitution on model membranes to evaluate the ability of protein-protein interactions, calcium, phosphorylation of the CFTR cytoplasmic domains, and increasing cholesterol density in membranes to promote CFTR cluster formation. Our computational model predicts that protein-protein interactions and membrane cholesterol density are key modulators of CFTR cluster formation. Testing of computational predictions on model membranes confirms that protein-protein interactions and increasing cholesterol density are essential for CFTR cluster formation. Phosphorylation of CFTR cytoplasmic regions and calcium modulate protein-protein interactions to control cluster formation. Increasing density of cholesterol in membranes results in increased CFTR cluster size and number. Furthermore, we find that CFTR clusters can merge on model membranes, suggesting that the condensed molecules are fluid. Combined, these data are consistent with properties seen for other membrane-associated phase-separated condensates [17,56–62]. These results strongly support phase separation as the driving mechanism for CFTR cluster formation.

## RESULTS

### Protein-protein interactions and membrane cholesterol are predicted to drive CFTR cluster formation *in silico*

Previous work by our lab [31,32] and others [35,37,49] demonstrated that the R region and C-terminal tail of CFTR can interact with neighboring R region and C-terminal tails and engage in multivalent interactions with NHERF-1, calmodulin, the sulphate transporter and anti-sigma factor antagonist (STAS) domains, including their disordered intervening sequence (IVS) regions and C-terminal tails of SLC26 family transporters; the transmembrane domains of CFTR can also interact with cholesterol in the membrane (Figure 1). We have previously demonstrated that multivalent interactions between membrane-associated proteins, binding partners, and cholesterol-rich membrane domains contribute to the formation of phase-separated biomolecular condensates [17,57–59]. Therefore, we posited that self-association between neighboring CFTR channels and multivalent interactions between CFTR and its binding partners, coupled to CFTR localization to cholesterol-rich membrane domains, could drive the formation of macromolecular clusters of CFTR [48–50]. To initially test our hypothesis, we created a computational biochemical model using Network Free Simulations (NFSIM) in the Virtual Cell modeling environment [63–65], which has previously been used to investigate phase separation between multivalent proteins [58,66–70]. Here we explicitly modeled experimentally demonstrated protein-protein and protein-cholesterol interactions for CFTR, calmodulin, NHERF-1, the SLC26A9 chloride transporter STAS domain and C-terminal tail (hereafter SLC26A9 STAS IVS-Cterm), and cholesterol (Figures S1, S2). These interactions include each of the N- and C-terminal lobes of calmodulin binding one of three sites in the R region of CFTR [32], and PDZ1 of NHERF-1 binding cholesterol localized in the membrane [33]. On (K_on_) and off (K_off_) rates for bimolecular interactions were taken directly from the literature, estimated from the literature (specifically lipid-protein rate constants), or experimentally derived by biolayer interferometry (BLI) using purified proteins (Figures 2A-D, Table 1). While these estimates and, in some cases, experimentally fit values are not precise, the exact numerical values are not critical for defining the qualitative behavior of our computational model. In this computational model, we also varied the density of CFTR and cholesterol on the membrane and the concentrations of soluble binding partners to test the importance of these parameters for CFTR cluster formation (Table 2).

**Figure 2.**
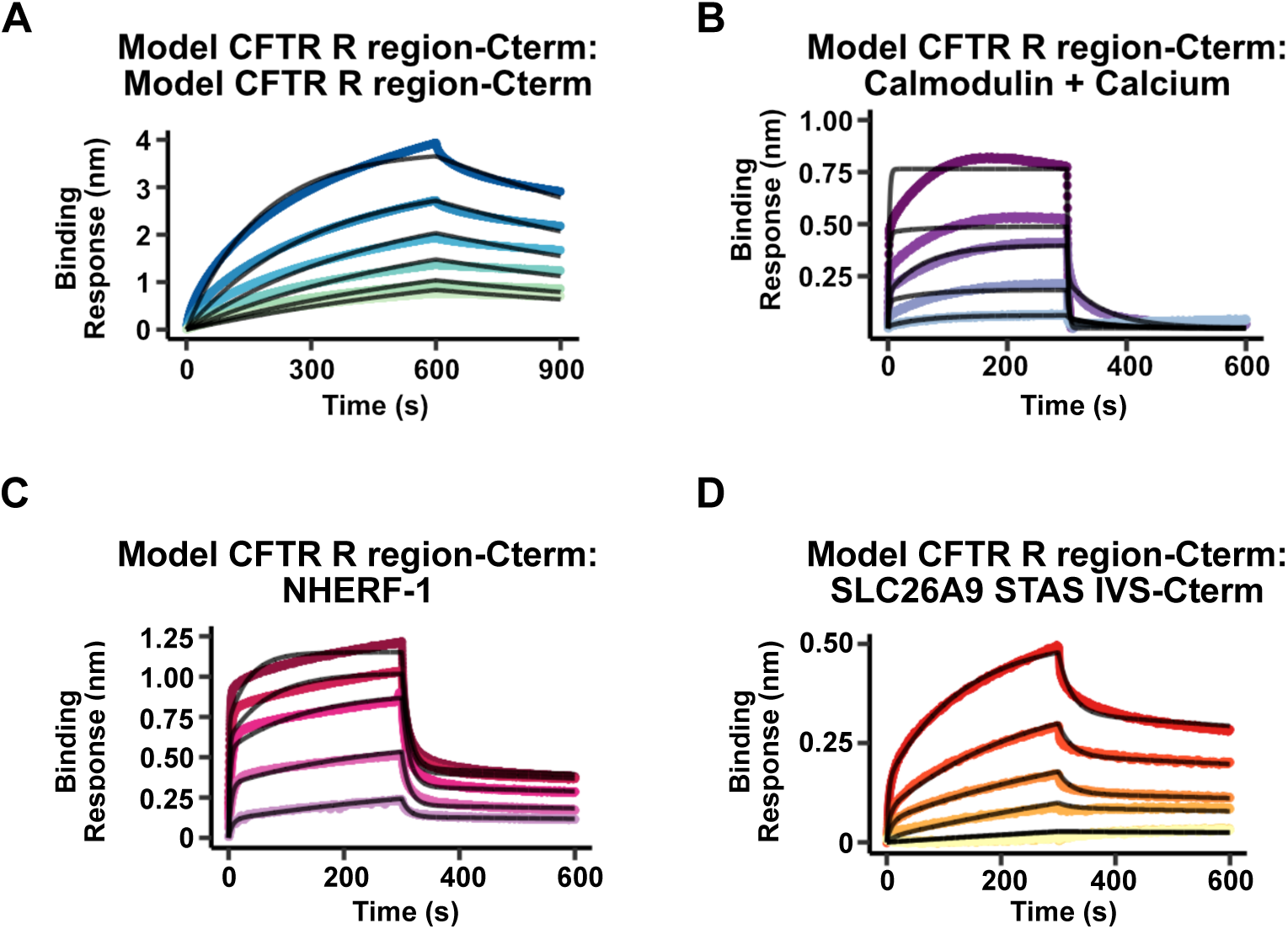
Biolayer interferometry binding curves. For **A-D**, colored lines are experimentally measured binding, black traces are modeled fits. **A)** Binding of Model CFTR R region-Cterm with itself. Model CFTR R region-Cterm was incubated in solutions diluted 2-fold from the initial solution that contained 25 μM Model CFTR R region-Cterm. **B)** Model CFTR R region-Cterm and calmodulin with calcium. Model CFTR R region-Cterm was incubated in solutions diluted 2-fold from the initial solution that contained 7.5 μM calmodulin. 500 μM CaCl_2_ was included in all solutions to ensure calcium-loaded calmodulin lobes. Note that the fits to the curves do not superimpose well with the data, reflecting a more complex binding mechanism of each of the two calmodulin lobes to at least three elements of the R region than our simple model with only two binding parameters. **C)** Model CFTR R region-Cterm and NHERF-1. Model CFTR R region-Cterm was incubated in solutions diluted 2-fold from the initial solution that contained 10 μM NHERF-1. **D)** Model CFTR R region-Cterm and SLC26A9 STAS IVS-Cterm. Model CFTR R region-Cterm was incubated in solutions diluted 2-fold from the initial solution that contained 10 μM SLC26A9 STAS IVS-Cterm. A total of 5 or 6 concentrations were used per analyte.

**Table 1.**
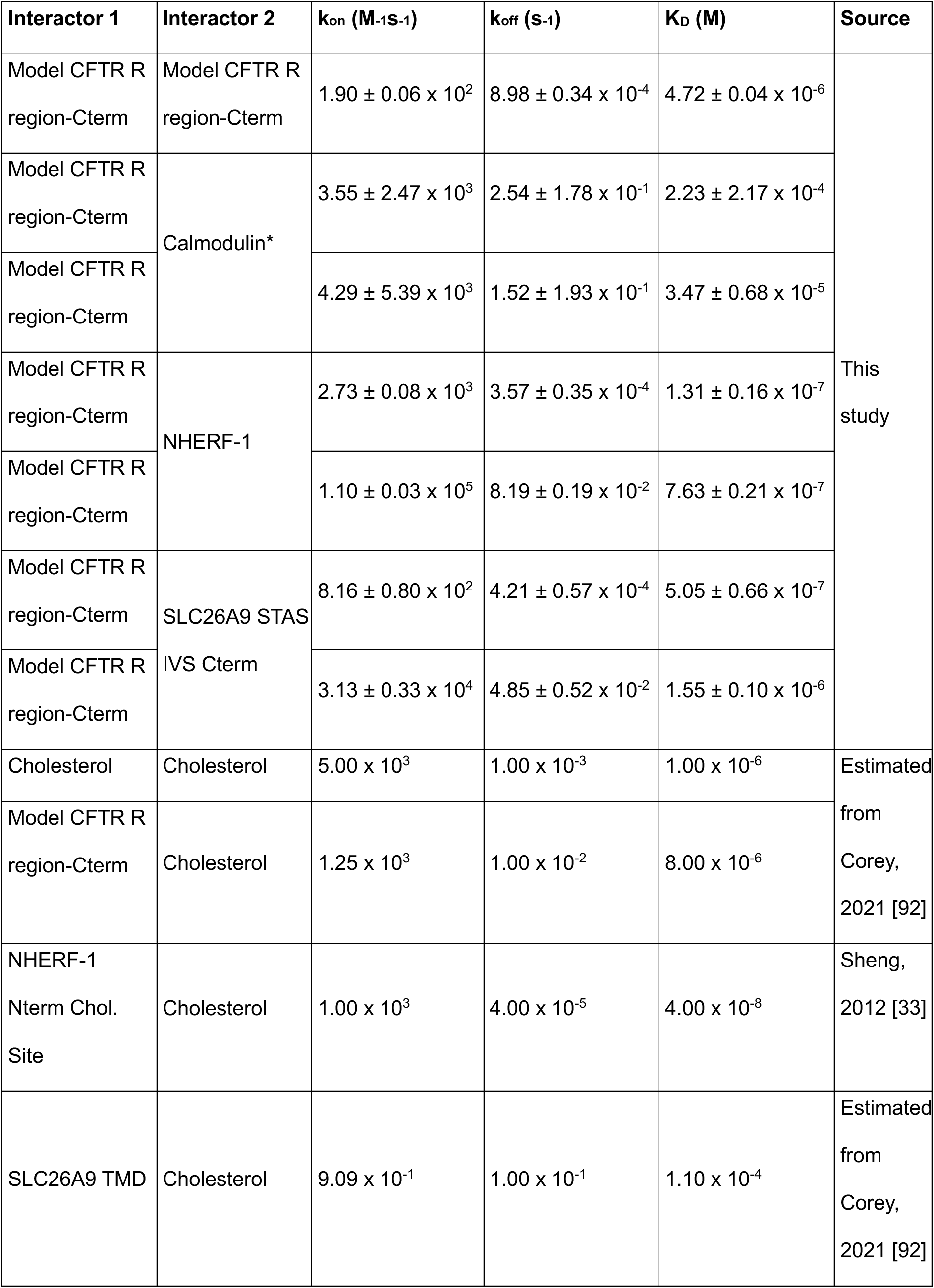
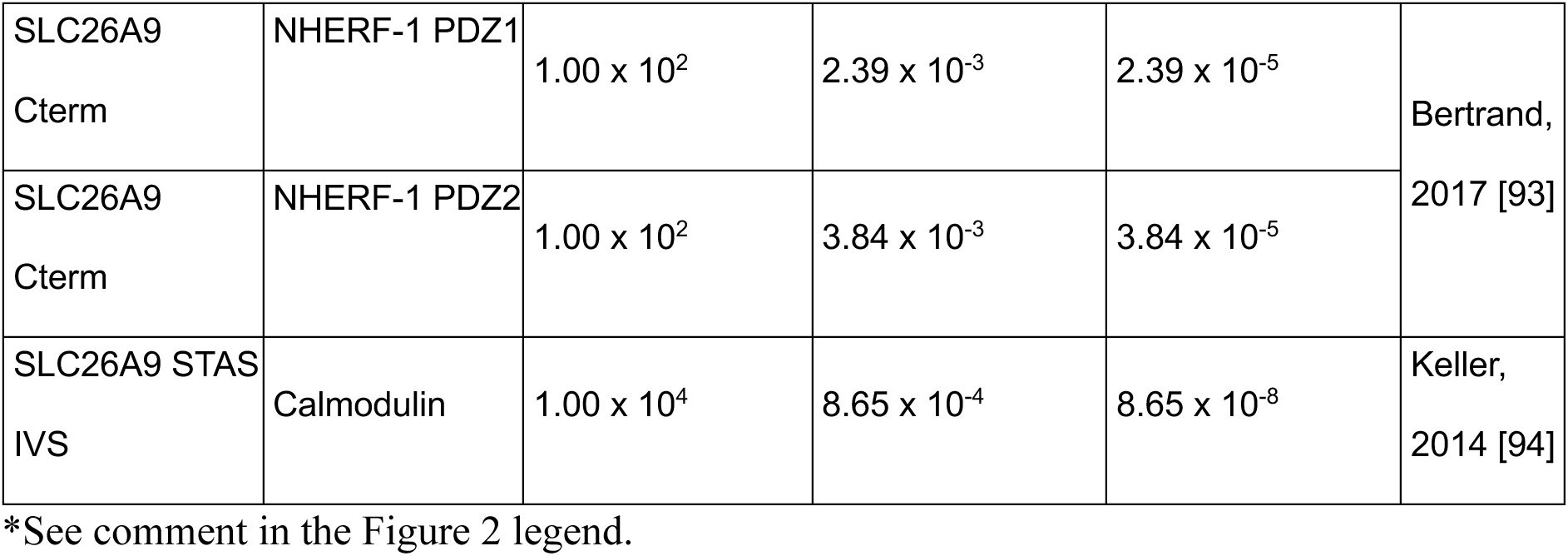
Binding constants used to describe interactions in the Virtual Cell computational model, based on Biolayer Interferometry experiments or literature values.

**Table 2.**
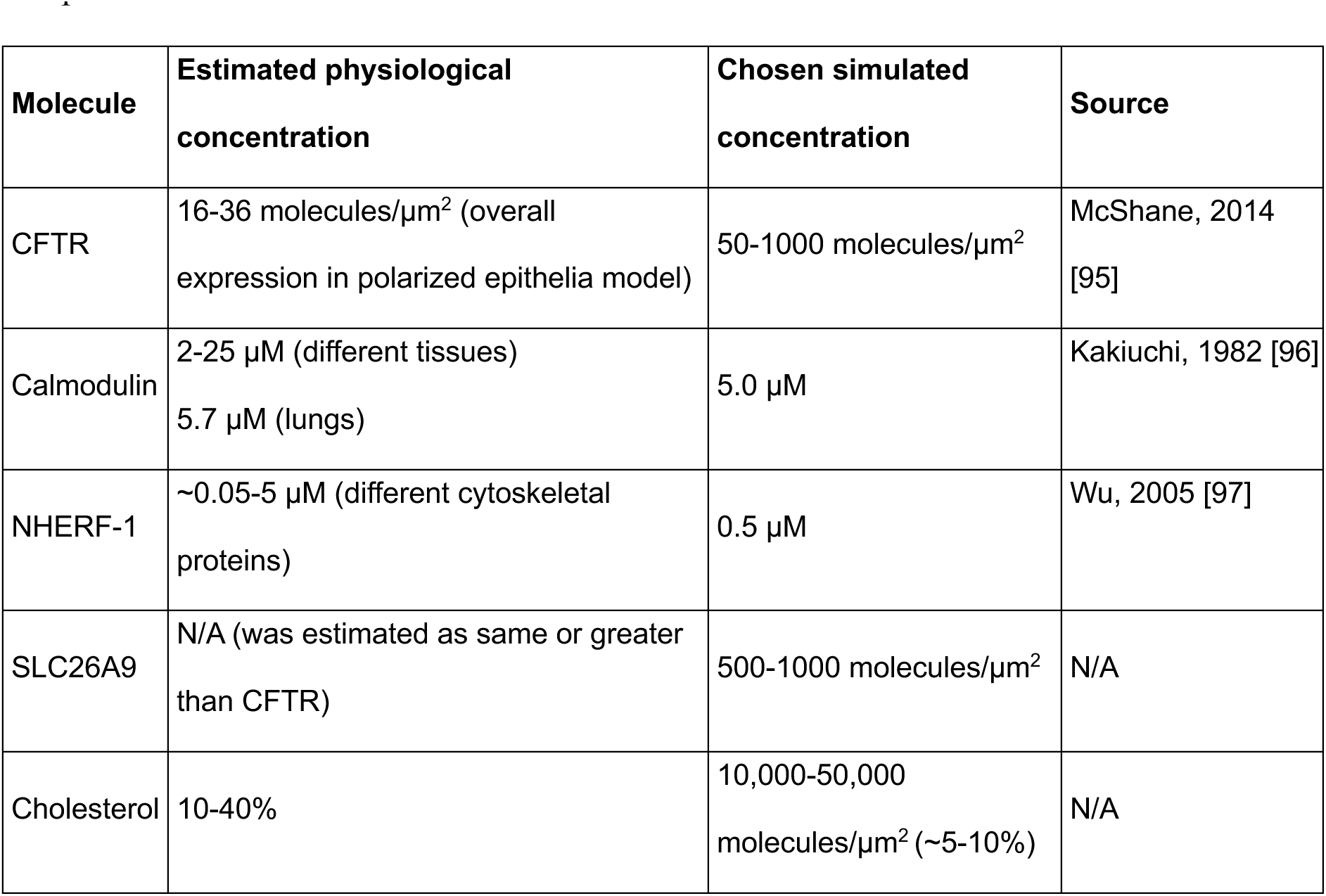
Densities and concentrations of molecular species used in the Virtual Cell computational model.

BLI measurements were performed using the following combinations of proteins: 1) CFTR R region and C-terminal tail (hereafter CFTR R region-Cterm) and CFTR R region-Cterm, 2) CFTR R region-Cterm and calcium-loaded calmodulin, 3) CFTR R region-Cterm and NHERF-1, and 4) CFTR R region-Cterm and SLC29A9 STAS IVS-Cterm. The binding curves for CFTR R region-Cterm self-association were best fit to a 1:1 bimolecular binding model with a binding constant of 4.72 µM (Figure 2A). The binding curves for CFTR R region-Cterm and calcium-loaded calmodulin were best fit to a 2:1 binding model with two binding constants of 34.7 µM and 223 µM (Figure 2B). The best fits are in agreement with NMR evidence that calmodulin binds separately to the CFTR R region via two distinct sites in the N-and C-terminal lobes of calmodulin; however, they are still not strongly overlapping the experimental data, likely reflecting the additional complexity of the binding to at least three R region elements revealed by NMR [32]. The binding curves for CFTR R region-Cterm and NHERF-1 were best fit to a 2:1 binding model that accounts for each NHERF-1 PDZ binding a single PDZ binding motif at the C-terminus of CFTR R region-Cterm with two binding constants of 0.131 µM and 0.763 µM (Figure 2C). NHERF-1 contains two PDZ domains that can each bind the PDZ binding motif found at the Cterm of CFTR. The two binding constants are likely correlated with the affinities of each NHERF-1 PDZ domain for CFTR R region-Cterm. Finally, the binding curves for CFTR R region-Cterm and SLC26A9 STAS IVS-Cterm were measured. Unlike calmodulin and NHERF-1, the specific interactions between the intrinsically disordered CFTR R region-Cterm and intrinsically disordered SLC26A9 STAS IVS-Cterm are not well characterized. However, the best fit for these binding curves was generated using a 2:1 binding model with two binding constants of 0.505 µM and 1.55 µM (Figure 2D). These binding constants were incorporated into our Virtual Cell computational model.

Using our Virtual Cell model, we first interrogated the ability of CFTR self-association and protein-protein interactions between CFTR R region-Cterm, SLC26A9 STAS IVS-Cterm, NHERF-1, and calmodulin, to drive the formation of CFTR clusters. Our model predicted that clusters containing 2 to 4 CFTR molecules could form on membranes as the density of CFTR on the membrane increased while binding partner concentrations were kept constant. No clusters containing more than 5 molecules of CFTR were predicted to form through self-association and protein-protein interactions alone, using our specified set of binding partners (Figure 3A). When we changed the interactions in our model to account for only CFTR self-association and cholesterol in membranes, our model predicted the formation of a single cluster containing greater than 100 molecules of CFTR in addition to the formation of smaller clusters containing more than 10, 5, or 2, CFTR molecules (Figure 3B). However, when we allowed for protein-protein and protein-cholesterol interactions, three clusters formed that contained more than 100 molecules of CFTR at the highest CFTR density used in the simulations (Figure 3C), dozens of clusters that contained more than 10 CFTR molecules at the two highest densities of CFTR formed, and tens to hundreds of clusters containing more than 5 or 2 molecules of CFTR formed (Figure 3C). Thus, our model predicts that CFTR cluster formation depends on specific protein-protein interactions, protein-lipid interactions, and the density of CFTR on membranes; these predictions are consistent with phase-separated condensate formation.

**Figure 3.**
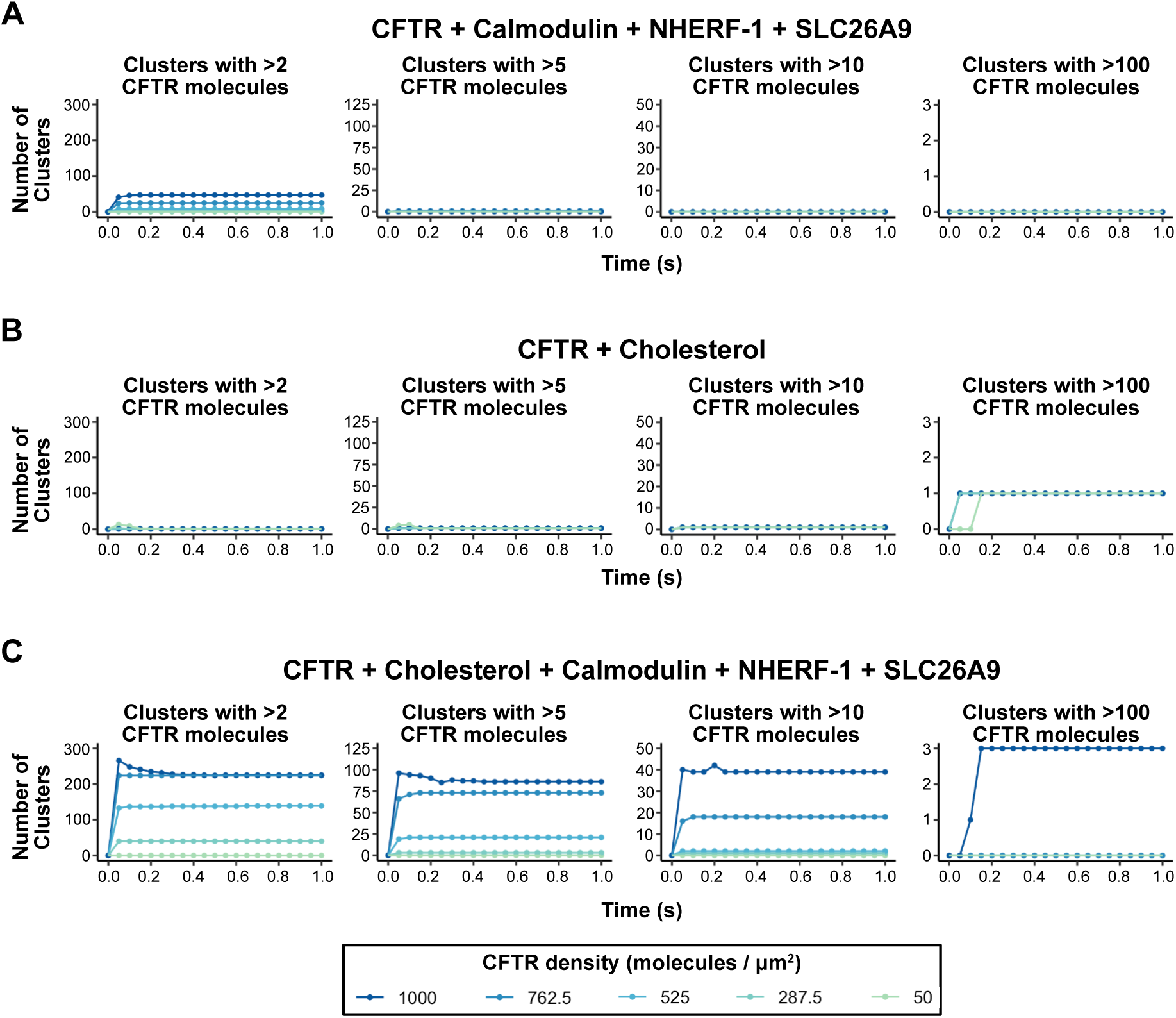
Simulations using the Virtual Cell modeling environment predict that protein – protein and membrane cholesterol – protein interactions are required for the formation of mesoscale CFTR-containing macromolecular clusters on membranes. Simulation results for the formation of clusters containing >2, >5, >10, or greater the >100 CFTR molecules for **A)** when only protein - protein interactions are considered, **B)** when only CFTR – membrane cholesterol interactions are considered, and **C)** when protein – protein and protein – membrane cholesterol interactions are considered.

### Wild type CFTR R region-Cterm self-associates on model membranes

We previously used *in vitro* supported lipid bilayers (SLBs) to probe the functional consequences of phase separation of T cell signaling proteins [57,59,71]. To study the intracellular disordered interacting element of CFTR, which had not previously been used on SLBs, we generated a single His_8_-tagged fusion of the CFTR R region and C-terminal region (hereafter sHis_8_-CFTR R region-Cterm) that can be attached to SLBs doped with NiNTA-modified lipids. We first tested whether AlexaFluor (AF) 488-labelled sHis_8_-CFTR R region-Cterm attached to the SLB specifically via NiNTA – histidine interactions or if AF488-sHis_8_-CFTR R region-Cterm associated with the membrane in an unexpected manner. We reasoned that if sHis_8_-CFTR R region-Cterm specifically associated with NiNTA-doped SLBs, high concentrations of imidazole would interrupt the interaction and AF488-sHis_8_-CFTR R region-Cterm would no longer associate with the membrane. We attached AF488-sHis_8_-CFTR R region-Cterm to SLBs and added buffer containing 0 mM or 275 mM imidazole. In buffer containing 0 mM imidazole, AF488-(s)His_8_-CFTR R region-Cterm remained attached to the membrane (Figure S3A, left) while additional of 275 mM imidazole resulted in a loss of AF488 fluorescence on the membrane (Figure S3A, right), indicating that its association was a direct result of NiNTA-modified lipid – histidine interactions and that there are no significant direct interactions of the membrane lipids with the CFTR R region or Cterm.

Membranes composed of complex lipid mixtures often undergo a liquid to solid transition when in direct contact with a glass coverslip substrate (Figure S3B) [17,72]. Therefore, in order to test our model predictions and experimentally probe the driving forces for CFTR cluster formation, we developed a SLB model in which we first laid a cholesterol-depleted SLB followed by a more complex, cholesterol-rich SLB, similar to those previous described for studying the biophysical properties of membranes [73]. We refer to this model membrane as a double supported lipid bilayer ((d)SLB). By protecting the complex lipid mixture of the upper bilayer from interacting with the coverglass substrate, we were able to maintain fluidity across a wide range of compositions (Figure S3C) (Table 3).

**Table 3.** Compositions of supported lipid bilayers. POPC is 1-palmitoyl-2-oleoyl-sn-glycero-3-phosphocholine, Avanti Cat. #850457. DPPC is 1,2-dipalmitoyl-sn-glycero-3-phosphocholine, Avanti Cat. #850355. Cholesterol was purchased from Avanti Cat. #700100. 18:1 DGS-NTA is 1,2-dioleoyl-sn-glycero-3-[(N-(5-amino-1-carboxypentyl)iminodiacetic acid)succinyl] (nickel salt), Avanti Cat. #790404. 16:0 DGS-NTA is 1,2-dipalmitoyl-sn-glycero-3-[(N-(5-amino-1-carboxypentyl)iminodiacetic acid)succinyl] (nickel salt), provided by the Levental Lab. PEG5000-PE is 1,2-dioleoyl-sn-glycero-3-phosphoethanolamine-N-[methoxy(polyethylene glycol)-5000] (ammonium salt), Avanti Cat. #880230.

| Figure | Bilayer | POPC | DPPC | Cholesterol | 18:1 DGS-NTA(Ni) | 16:0 DGS-NTA(Ni) | PEG5000PE |
| --- | --- | --- | --- | --- | --- | --- | --- |
| 5, 6, 7 | Base | 99% | 0% | 0% | 0% | 0% | 1% |
| 4 | Base | 98% | 0% | 0% | 2% | 0% | 0.01% |
| 5 | Top | 59% | 29% | 10% | 2% | 0% | 0.01% |
| 5, 6 | Top | 59% | 29% | 10% | 0% | 2% | 0.01% |
| 5 | Top | 49% | 29% | 20% | 2% | 0% | 0.01% |
| 5 | Top | 49% | 29% | 20% | 0% | 2% | 0.01% |
| 5 | Top | 39% | 29% | 30% | 2% | 0% | 0.01% |
| 5 | Top | 39% | 29% | 30% | 0% | 2% | 0.01% |
| 5 | Top | 29% | 29% | 40% | 2% | 0% | 0.01% |
| 5, 6, 7 | Top | 29% | 29% | 40% | 0% | 2% | 0.01% |

In our previous NMR spectroscopy studies using a non-His-tagged version of this fusion protein [31,32], we noted that the R region was prone to aggregation in solution. We generated a double-His_8_-tagged version of the disordered CFTR intracellular R region and C-terminal tail (hereafter WT CFTR R Region-Cterm, Figure S4) that enables NiNTA-modified lipid interactions at both the N-terminus and between the R region and C-terminal elements, better mimicking the topology of the full-length membrane protein. To test whether this fusion protein formed clusters on membranes without the contribution of other interactions, we attached WT CFTR R region-Cterm via interactions of the two His_8_ tags to NiNTA-modified lipids doped in SLBs at 2 mol%. We immediately observed small clusters of WT CFTR R region-Cterm on the membrane, as indicated by increased appearance of puncta in each image, across a wide range of WT CFTR R Region-Cterm concentrations (Figure 4A). Fluorescence recovery after photobleaching (FRAP) analysis of non-clustered regions showed near full recovery of fluorescence, indicating that most WT CFTR R Region-Cterm molecules are mobile on the bilayer (Figure 4B), even with the evidence for self-association.

**Figure 4.**
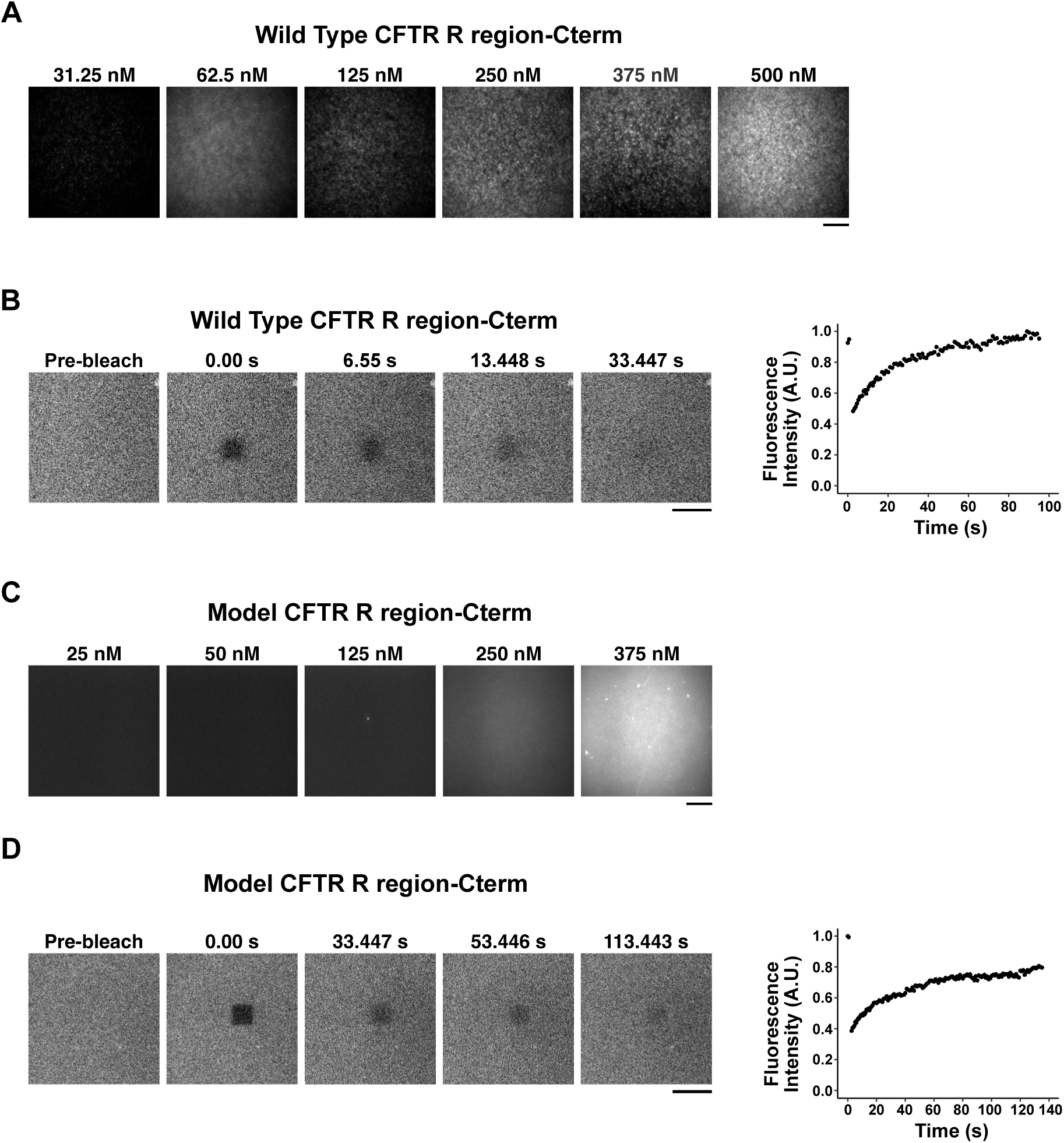
Model CFTR R Region-Cterm is evenly distributed on supported lipid bilayers (SLBs) except at the highest concentration. **A)** AF647-WT CFTR R region-Cterm fluorescence is not evenly distributed on SLBs as the density increases (left), as indicated by the variance in the plot (right). **B)** Fluorescence recovery after photobleaching (FRAP) indicates that AF647-WT CFTR R Region-Cterm is fluid on SLBs. **C)** AF647-Model CFTR R region-Cterm fluorescence is evenly distributed on SLBs as the density increases. **D)** Fluorescence recovery after photobleaching (FRAP) indicates that AF647-model CFTR R Region-Cterm is fluid on SLBs. Scale bars = 30 µm

Because we could not quantify the contribution of WT CFTR R region-Cterm to the observed cluster formation, we sought to generate a model CFTR R region-Cterm lacking self-associating regions that could be used to investigate other protein-protein and protein-lipid interactions contributing to CFTR cluster formation. To create this fusion protein, we returned to our previous NMR data [31,32] and retained only regions having interactions with binding partners in our double His_8_-tagged fusion (hereafter Model CFTR R region-Cterm, see Materials and Methods, Figure S4). We attached the Model CFTR R region-Cterm to (d)SLBs via NiNTA-modified lipid interactions to the two His_8_ tags and observed homogenously distributed fluorescence across the membrane as the concentration of Model CFTR R Region-Cterm was increased (Figure 4C). Model CFTR R Region-Cterm attached to the (d)SLB also exhibited near-full recovery of fluorescence following photobleaching (Figure 4D). Combined, these results indicate that WT CFTR R region-Cterm exhibits self-association on (d)SLBs. Model CFTR R region-Cterm is well distributed on the (d)SLB and lacks the propensity to self-associate to a high enough degree to observe resolvable clusters. Therefore, we deemed the Model CFTR R region-Cterm fusion protein to be most suitable for subsequent reconstitution experiments, since changes to protein distribution and dynamics as a result of changing experimental conditions can be readily assessed.

### NHERF-1, SLC26A9 STAS IVS-Cterm, calmodulin, and calcium are insufficient for inducing CFTR R region-Cterm cluster formation *in vitro*

Previous work using NMR demonstrated that calcium enhances calmodulin N- and C-terminal lobe binding to the CFTR R region [32]. We also observed that calcium promotes binding of calmodulin with the CFTR R region using BLI (Figures 2B). However, our computational model predicts that only small, sub-resolution (clusters containing less than 5 molecules of CFTR would not be resolvable using fluorescence microscopy) CFTR R region-Cterm clusters will form upon addition of binding partners, including calcium-loaded calmodulin. Given our previous NMR and current BLI data (Figures 2A-D, Table 1), the reported link between CFTR and calcium signaling [50], calcium-calmodulin mediated activation of CFTR [32], and our model predictions, we were highly interested in experimentally interrogating the role of protein-protein interactions including calcium-loaded calmodulin in driving CFTR clustering. We first tested the ability of a combination of 0.5 µM NHERF-1, 1 µM SLC26A9 STAS IVS-Cterm, and 1 µM calmodulin with and without 500 µM CaCl_2_ to induce visible cluster formation of Model CFTR R region-Cterm. The addition of this mixture did not result in visible Model CFTR R region-Cterm cluster formation (Figure 5A, left column). Our experimental results are consistent with our computational predictions: protein-protein interactions among our set of CFTR binding partners are not sufficient to drive the formation of visible clusters of Model CFTR R region-Cterm using these protein concentrations.

**Figure 5.**
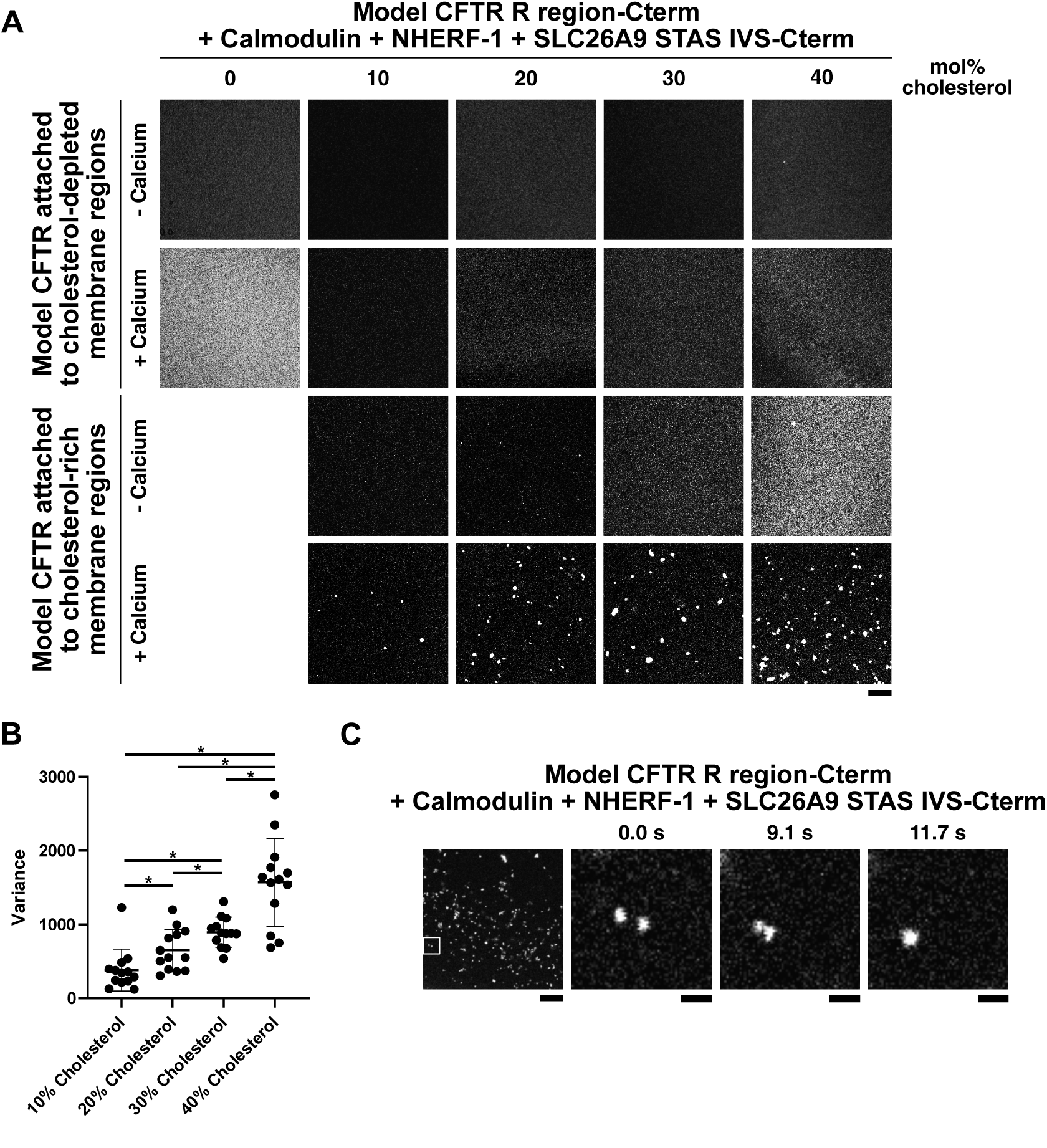
Cholesterol, binding partners, and calcium promote model CFTR R region-Cterm cluster formation on double supported lipid bilayers ((d)SLBs). **A)** Model CFTR R region-Cterm attached to cholesterol-depleted regions of (d)SLBs do not form clusters in the presence of 1 µM calmodulin, 0.5 µM NHERF-1, and 1 µM SLC26A9 STAS IVS-Cterm without or with 500 µM CaCl_2_ (top two rows). Model CFTR R Region-Cterm attached to cholesterol-rich regions of the (d)SLB forms clusters in the presence of 1 µM calmodulin, 0.5 µM NHERF-1, and 1 µM SLC26A9 STAS IVS-Cterm with 500 µM CaCl_2_ (bottom row) but not without 500 µM CaCl_2_ (third row). Scale bar = 30 µm. **B)** Variance of fluorescence intensities in **(A)** demonstrates significantly increased CFTR R region-Cterm cluster formation as the density of cholesterol is increased in (d)SLBs. n = 13. * = p < 0.05, T-test. **C)** Full field of view of Model CFTR R region-Cterm clusters (left). Scale bar is 15 µm. Time-lapse imaging of Model CFTR R region-Cterm clusters shows merging event between Model CFTR R region-Cterm clusters (images expanded from box inset in left image). Scale bar = 5 µm.

### Cholesterol and protein-protein interactions drive the formation of Model CFTR R region-Cterm clusters *in vitro*

Because our computational model predicts that cholesterol interactions with CFTR are necessary for robust CFTR cluster formation, we investigated the role of cholesterol in modulating CFTR clustering *in vitro.* We initially performed experiments with Model CFTR R region-Cterm attached to 18:1 DGS-NTA nickel modified lipids localized to cholesterol-depleted regions of (d)SLBs (Figure S5A) with 0.5 µM NHERF-1, 1 µM SLC26A9 STAS IVS-Cterm, and 1 µM calmodulin with and without 500 µM CaCl_2_ in solution. Visible clusters of Model CFTR R region-Cterm did not form as the cholesterol density was increased from 0 to 40 mol% (Figure 5A, top two rows). We then repeated these experiments with Model CFTR R Region-Cterm attached to 16:0 DGS-NTA localized in cholesterol-rich regions of (d)SLBs (Figure S5B). We previously used 16:0 DGS-NTA to localize His-tagged proteins to cholesterol-rich membrane domains in spin-coated multi-layered model bilayers [17]. We first performed experiments using (d)SLBs composed of increasing cholesterol density from 0 to 40 mol%. We attached Model CFTR to cholesterol-rich (d)SLB regions and incubated with 0.5 µM NHERF-1, 1 µM SLC26A9 STAS IVS-Cterm, and 1 µM calmodulin without including CaCl_2_ in solution. We did not observe robust cluster formation, although sparse mesoscale clusters did appear when (d)SLBs contained 40 mol% cholesterol (Figure 5A, third row). However, upon addition of CFTR R region-Cterm binding partners and 500 µM CaCl_2_, we observed the formation of Model CFTR R region-Cterm clusters (Figure 5A, bottom row). Cholesterol had a dose dependent effect on clustering, where cluster number and area increased non-linearly with increases in cholesterol (Figure 5B). At 10 and 20 mol% cholesterol, clusters were small (2-3 µm^2^), and generally spherical. At 30 and 40 mol% cholesterol, clusters were larger (>3 µm^2^) and more irregular in shape. Clusters across all cholesterol densities tested appeared dynamic, as multiple fusion events of two small clusters merging into a single large cluster were observed (Figure 5C). These results are consistent with predictions from our computational model.

Using our (d)SLB experimental system, we then sought to understand which specific protein-protein interactions were important for Model CFTR R region-Cterm cluster formation. We formed (d)SLBs containing either 10 or 40 mol% cholesterol and attached Model CFTR R region-Cterm to cholesterol-rich regions. We then added 0.5 µM NHERF-1 and / or 1 µM SLC26A9 STAS IVS-Cterm to solutions containing 1 µM calmodulin and 500 µM calcium. We observed no Model CFTR R region-Cterm cluster formation when either NHERF-1 or SLC26A9 STAS IVS-Cterm was present in the solution with calmodulin and calcium (Figure 6A). However, when both were present in solution with calmodulin and calcium, clusters of Model CFTR R region-Cterm formed on membranes composed of 10 or 40 mol% cholesterol, with a higher number and larger size of clusters on membranes composed of 40 mol% cholesterol (Figure 6A). Experiments were repeated, this time using calmodulin and calcium as the Model CFTR R region-Cterm binding partners with 0.5 µM NHERF-1 and 1 µM SLC26A9 STAS IVS-Cterm already in solution. Neither 1 µM calcium alone nor 500 µM calmodulin induces Model CFTR R region-Cterm cluster formation on (d)SLBs in the presence of NHERF-1 and SLC26A9 STAS IVS-Cterm (Figure 6B). When both were combined in solution, small sparse Model CFTR R region-Cterm clusters formed on (d)SLBs composed of 10 mol% cholesterol while large more numerous clusters formed on (d)SLBs composed of 40 mol% cholesterol (Figure 6B). These results demonstrate that observable Model CFTR R region-Cterm cluster formation is induced only in the presence of all tested components; lack of any component inhibits Model CFTR R region-Cterm cluster formation.

**Figure 6.**
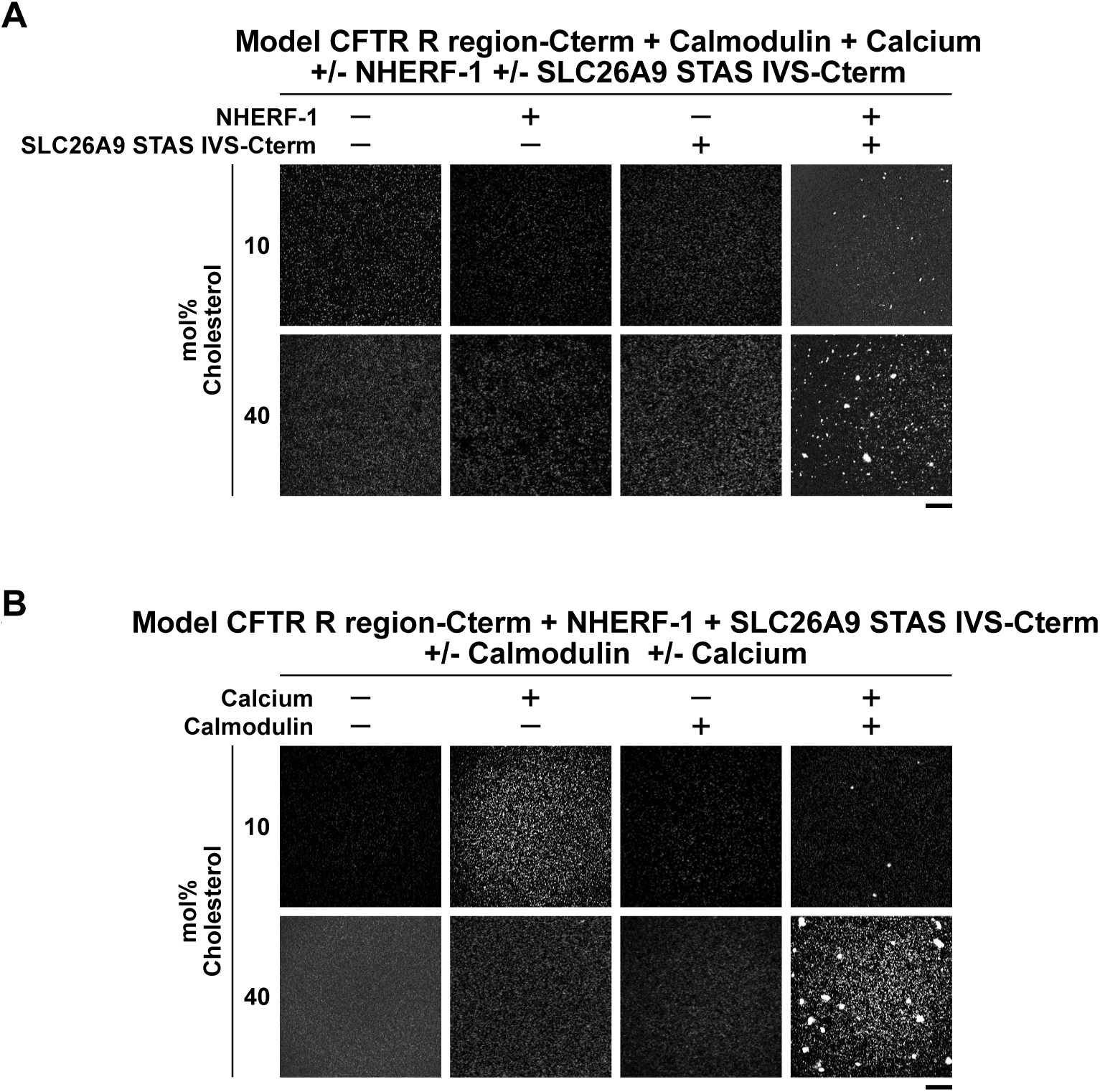
Distinct sets of CFTR R region-Cterm binding partners promote cluster formation on double supported lipid bilayers ((d)SLBs). **A)** 0.5 µM NHERF-1 and 1 µM SLC26A9 STAS IVS-Cterm promote cluster formation of AF647-Model CFTR R region-Cterm attached to cholesterol-rich (d)SLB regions when combined in solution. Scale bar = 30 µm. **B)** 1 µM calmodulin and 500 µM CaCl_2_ promote cluster formation of AF647-Model CFTR R region-Cterm attached to cholesterol-rich (d)SLB regions when combined in solution. Scale bar = 30 µm. Proteins in **(A)** and **(B)** were incubated for 30 minutes prior to image capture.

### Phosphorylation of the R region alters membrane distribution of Model CFTR R region-Cterm clusters *in vitro*

PKA-mediated phosphorylation is a mechanism of CFTR channel activation that acts as an alternative to calcium-calmodulin activation [32]. Given these opposing functions, we were interested in the impact of CFTR R region-Cterm phosphorylation on clustering. We phosphorylated Model CFTR R region-Cterm using PKA and attached the non-phosphorylated or phosphorylated protein onto (d)SLBs composed of 40 mol% cholesterol and incubated with 1 µM calmodulin, 0.5 µM NHERF-1, and 1 µM SLC26A9 STAS IVS-Cterm with and without 500 µM calcium (Figure 7A). Non-phosphorylated Model CFTR R region-Cterm forms micron-sized clusters only in the presence of calcium (Figure 7B). Phosphorylation induces the formation of smaller more numerous clusters of Model CFTR R region-Cterm clusters, when compared with those formed by non-phosphorylated Model CFTR R region-Cterm, independent of buffer calcium (Figure 7C). These results demonstrate that protein-protein interactions involving either calcium-calmodulin or phosphorylation of the cytoplasmic domains of CFTR can induce cluster formation when combined with cholesterol in membranes.

**Figure 7.**
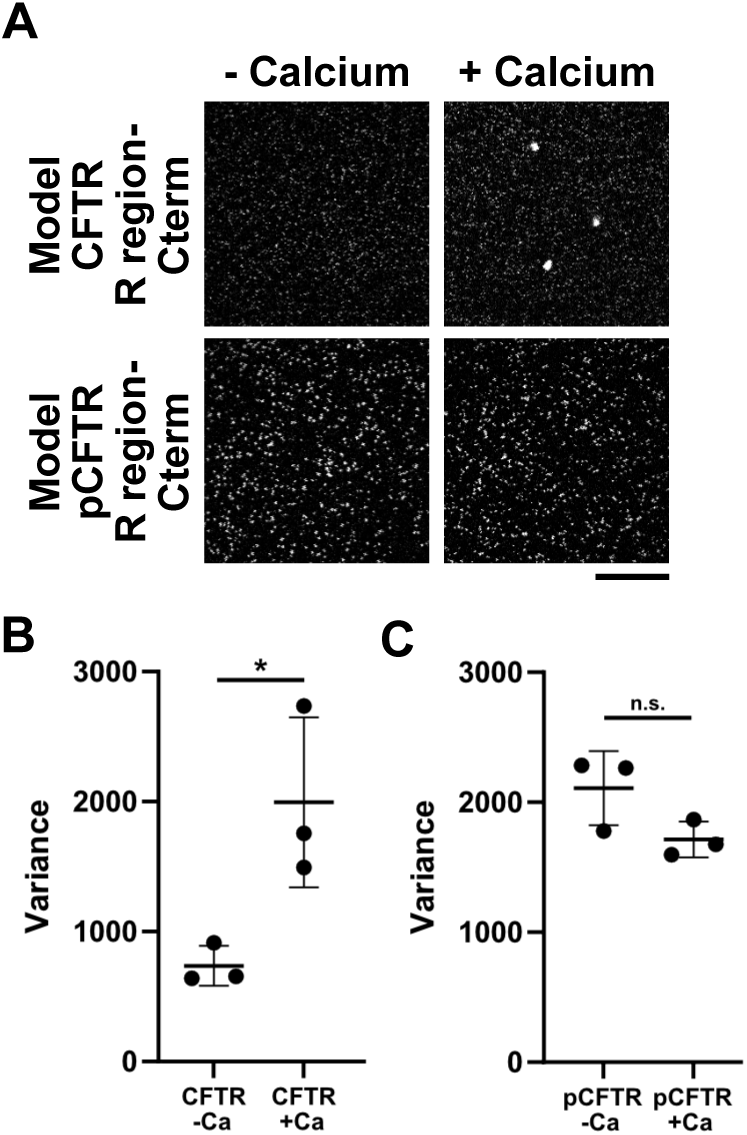
Phosphorylation of Model CFTR R region-Cterm results attached to cholesterol-rich regions of double supported lipid bilayers ((d)SLBs) results in cluster formation. **A)** AF647-Model CFTR R region-Cterm cluster formation on (d)SLBs is promoted in the presence of 1 µM calmodulin, 0.5 µM NHERF-1, and 1 µM SLC26A9 STAS IVS-Cterm with 500 µM CaCl_2_ (top row). Phosphorylation of AF647-Model CFTR R region-Cterm cluster promotes formation of smaller clusters on (d)SLBs in the presence of 1 µM calmodulin, 0.5 µM NHERF-1, and 1 µM SLC26A9 STAS IVS-Cterm with and without 500 µM CaCl_2_ (bottom row). Scale bar = 30 µm. **B)** Variance of AF647-Model CFTR R region-Cterm fluorescence intensities in the presence of calmodulin, NHERF-1, and SLC26A9 STAS IVS-Cterm with or without calcium. n = 3. * = p < 0.05, T-test. **C)** Variance of phosphorylated AF647-Model CFTR R region-Cterm fluorescence intensities in the presence of calmodulin, NHERF-1, and SLC26A9 STAS IVS-Cterm with or without calcium. n = 3. n.s. = no significance, p > 0.05, T-test.

## DISCUSSION

In this study, we demonstrate that CFTR cluster formation depends on multivalent protein-protein interactions, calcium, and cholesterol in the membrane, which is consistent with phase separation as the driving physical mechanism for CFTR cluster formation on membranes. CFTR clusters can merge with one another, indicating that the molecules within the clusters are dynamic. We found that phosphorylation of the cytoplasmic domains of CFTR can drive cluster formation with the full complement of binding partners and cholesterol, without calcium in the buffer (Figure 7). This observation may be partly explained by the propensity for the phosphorylated CFTR R region binding to the CFTR C-terminal tail. These results are complementary to previous reports that described interactions between phosphorylated R region with specific binding partners, including the C-terminal tail of CFTR, independent of non-phosphorylated CFTR R region interactions with calmodulin, and that phosphorylation of the R region by PKA inhibits calmodulin binding [32]. This potential for independent inputs into cluster formation is reminiscent of LAT with Grb2 and Sos1 or Gads and SLP-76 [57] or integrins with kindlin or talin in focal adhesions [60,74] and provides an example of the general principle of redundancy amongst membrane-associated protein interacting partners. Such redundancy could enable unique modes of CFTR cluster formation on the membranes of various cell types and may be especially important because not all types of cells express every CFTR interactor; ionocyte, ciliated, and basal cells in the airway epithelia express varied levels of CFTR and interactors [75,76].

Our study also generated an editable, user-friendly computational model housed in the Virtual Cell modeling environment. This tool is freely available to the community and can be adapted by researchers to investigate CFTR interactions with other binding partners and predict the formation of macromolecular clusters on membrane surfaces. Furthermore, the model can be adapted to account for phosphorylation and other posttranslational modifications to test their effect on protein-protein interactions and the subsequent role these modifications have on CFTR cluster formation. Additionally, the computational model can be used to predict the effect of small molecule therapeutics on interactions within CFTR clusters. This type of tool will be valuable for predicting off-target effects in CFTR interaction network behavior linked with small molecule interactions if the effects of the small molecule on binding affinities have been evaluated. Combined, our computational model and *in vitro* biochemical reconstitution system can be used to parse the complex co-regulation observed in cells to determine intracellular components that contribute to the co-clustering of CFTR and binding partners.

Finally, the results we present here enable the viewing of CFTR cluster formation through the lens of biological phase separation. The phase separation of membrane-associated proteins has been linked with T cell and B cell activation [17,18,57,59,61,62], glomerular filtration barrier maintenance [9,56,58], synaptic plasticity [77,78], cell adhesion [60,74], and growth factor receptor signaling pathways [79]. To date, channel and transporter function has yet to be linked with phase separation-directed organization. Here we provide evidence that CFTR can undergo phase separation with its multivalent binding partners coupled to cholesterol in the membrane. Importantly, wild type CFTR intracellular domains can self-associate to a high degree, thereby increasing the effective valency of interacting motifs within the R region and C-terminal tail when arrayed on membranes. In our study, we intentionally used a model CFTR fusion of the R region and C-terminal tail with a decreased propensity for CFTR cytoplasmic domain self-association in our experiments (Figure 4) so that we could specifically parse the contribution of binding partners and membrane components to CFTR cluster formation. In cells, self-association of CFTR will lower the density and concentration thresholds above which CFTR and its binding partners will undergo phase separation. Thus, clusters of CFTR that have been previously observed [48–50] likely form through phase separation akin to signaling clusters described above.

Another aspect of phase separation underlying the formation of CFTR clusters is the emergence of self-organized CFTR clusters with distinct size distribution profiles. These profiles depend on 1) the percent of cholesterol in the membrane or 2) the type of interactions that drive cluster formation. We observed that CFTR R region-Cterm localization with cholesterol was required for mesoscale cluster formation in our computational simulations and *in vitro* reconstitution experiments (Figures 3 and 5). Furthermore, as the percentage of cholesterol increased in model membranes, CFTR R region-Cterm clusters increased in size and number (Figure 5A). Together, these data show a direct role of membrane composition in the modulation of CFTR cluster formation, size, and number. When clustering is promoted by binding partner interactions, large clusters form (Figure 7A, top row). When clustering is promoted by interactions between phosphorylated CFTR R region-Cterm in the presence of binding partners, smaller more numerous clusters form relative to the binding partner alone condition (Figure 7A, bottom row). Importantly, the total amount of phase separation, as measured by the variance in fluorescence, is similar between CFTR R region-Cterm and binding partners or phosphorylated CFTR R region-Cterm and binding partners with or without calcium (Figure 7B, + calcium vs. Figure 7C, +/- calcium). This is especially pertinent, as a recent computational study shows that membrane-associated condensate function (specifically, actin polymerization) can be controlled by cluster size distribution [70]. The authors report that more numerous smaller clusters collectively promote a higher amount of actin polymerization than fewer, large clusters. Given that actin polymerization is increased at condensates because condensates increase the probability of Arp2/3 activation [58], and that CFTR open probability may be modulated by cluster formation, the possibility exists that cells may be able to modulate CFTR channel activity by promoting unique modes of cluster formation, i.e., with significant contribution from binding of R region to either calcium-calmodulin or the CFTR C-terminus in the presence of phosphorylation. Thus, CFTR cluster formation by phase separation may allow for tuning of CFTR function in different cell types or under different conditions by specifically regulating plasma membrane lipid composition, binding partner expression, phosphorylation of the intracellular domains of CFTR, and maintaining specific size distributions on the plasma membrane.

It is becoming increasingly evident that CF results in lipid imbalances in cellular membranes [80]. Several studies have shown that CFTR regulates lipid metabolism and homeostasis [81–84]. Furthermore, lipid interactions with CFTR and ENaC are required for optimal channel function [85]. This bi-directional relationship between lipids and CFTR is further highlighted by our results reported here, in which we show increased cluster formation directly linked with increased cholesterol density in model membranes (Figures 5 and 6). Therefore, it will be important for future studies to investigate specific mechanisms by which lipid components of membranes, including cholesterol and sphingolipids contribute to functional CFTR cluster formation, and the effect of CFTR clustering on lipid metabolism and homeostasis.

The results reported here are important for at least two reasons: 1) phase separation of membrane-associated proteins can be functionally coupled to lipid phase separation [17], and 2) manipulation of phase separation by novel therapeutics provides a new avenue for treating disease [86,87]. In cells, prevention of CFTR cluster formation is correlated with decreased CFTR channel function during vasoactive intestinal peptide stimulation [48]. However, the functional coupling between protein condensates and lipid domains for CFTR has yet to be shown. Combined, these data from our study and previous cellular data [48–50] lead us to propose a tantalizing possibility: CFTR cluster formation results in increased channel function per molecule, or, alternatively, in regulated channel function in a manner dependent on the components within the condensate, as has been reported for other membrane-associated and cytoplasmic phase-separating systems [11,57,58,62,88,89]. Thus, we suggest that phase separation of CFTR provides a possible mechanism for increased or otherwise regulated function of CFTR and other proteins co-localized in CFTR condensates on membranes, including effects on open channel probabilities. Future studies to test this hypothesis are needed, with the potential to enrich understanding of the regulation of CFTR and more generally of cholesterol-rich domain-localized membrane proteins, particularly those thought to be “co-regulated” with other membrane proteins.

## MATERIALS AND METHODS

### Modeling CFTR clustering with Virtual Cell

The Virtual Cell platform [63–65] was used to model and simulate CFTR cluster formation (https://vcell.org/). For each interaction modelled, binding mechanisms and kinetic constants were based either on biolayer interferometry data from this study or published data and are detailed in Table 1 provided in the supplementary material. Table 1 contains the rate constants and Table 2 contains the densities and concentrations of molecular species in the models. The models used in this study can be found in the Virtual Cell public database under user “ywan” with the following model name: “CFTR Membrane Clustering.”

### Expression constructs

Sequences of all constructs are included in Table 4 with schematics shown in Supplemental Figure S6A. CFTR R region-Cterm insert sequences were codon-optimized and ordered from GenScript. CFTR R region-Cterm gene inserts were amplified by PCR and cloned into pET plasmids using Gibson assembly. All R region sequences contain an F833L polymorphism that improves the solubility of the fusion proteins [31,32]. To generate the Model CFTR R Region-Cterm sequence, we used previously published NMR data acquired in our lab [31,32] and removed residues that were not found to have significantly broadened NMR resonances upon interaction with calmodulin or other protein binding partners (Figure S4). WT and Model CFTR R region-Cterm constructs were poly-histidine tagged with His_8_ for membrane attachment via nickel-chelated lipid anchors [56,57,71,90]. Full length calmodulin was inserted into a pET3a plasmid, while NHERF-1 (residues 1-358) and the intervening sequence and C-terminal region of SLC29A9 (STAS IVS-Cterm, residues 569-652, 742-772, and 785-791 in a single, fused disordered chain) were inserted into pET-His-SUMO plasmids. All inserts were validated using Sanger sequencing. Plasmids were transformed into, amplified in, and purified from NEB 10-beta Competent *E. coli* cells using the QIAprep Miniprep kit according to the manufacturer’s instructions.

**Table 4.**
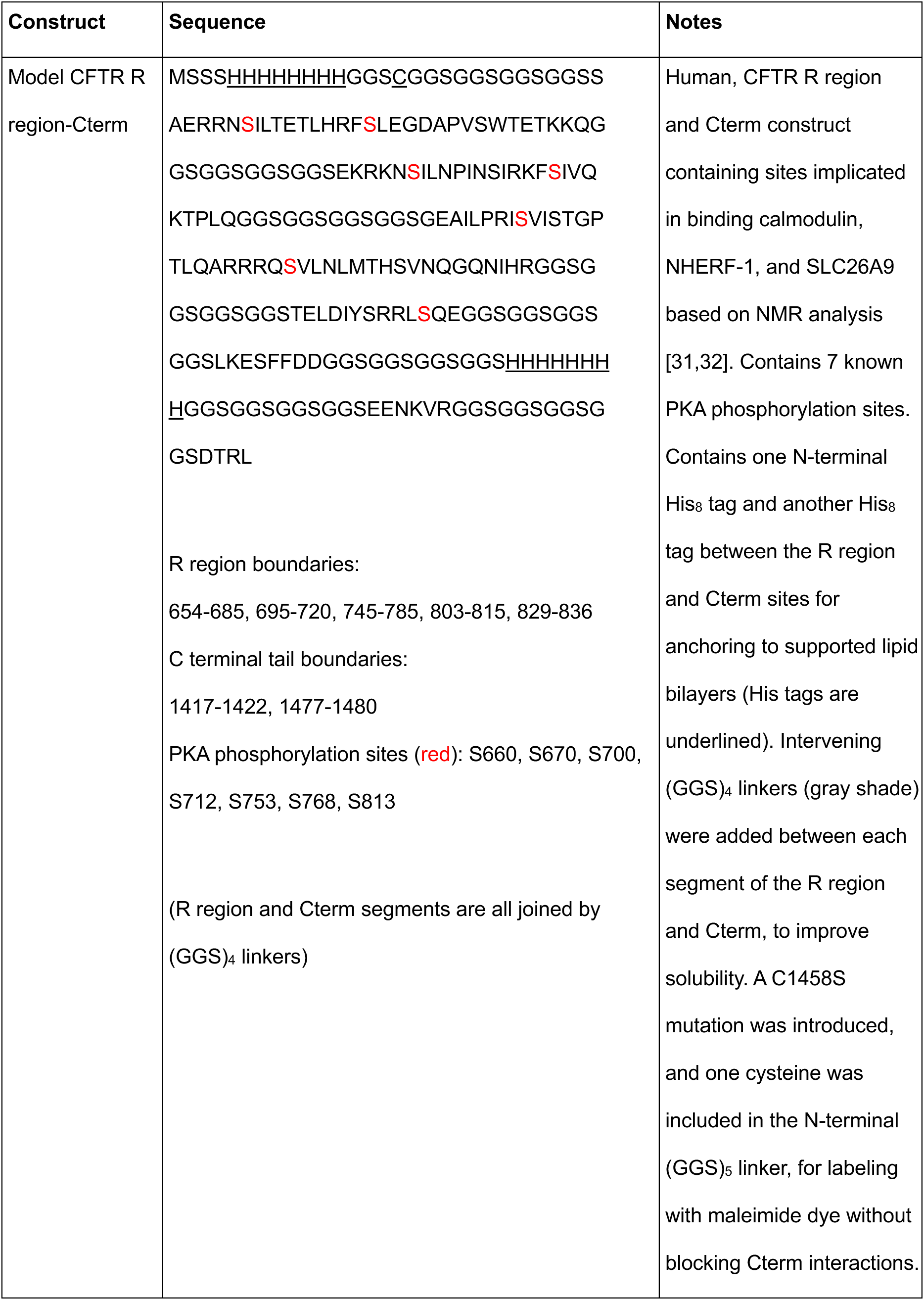
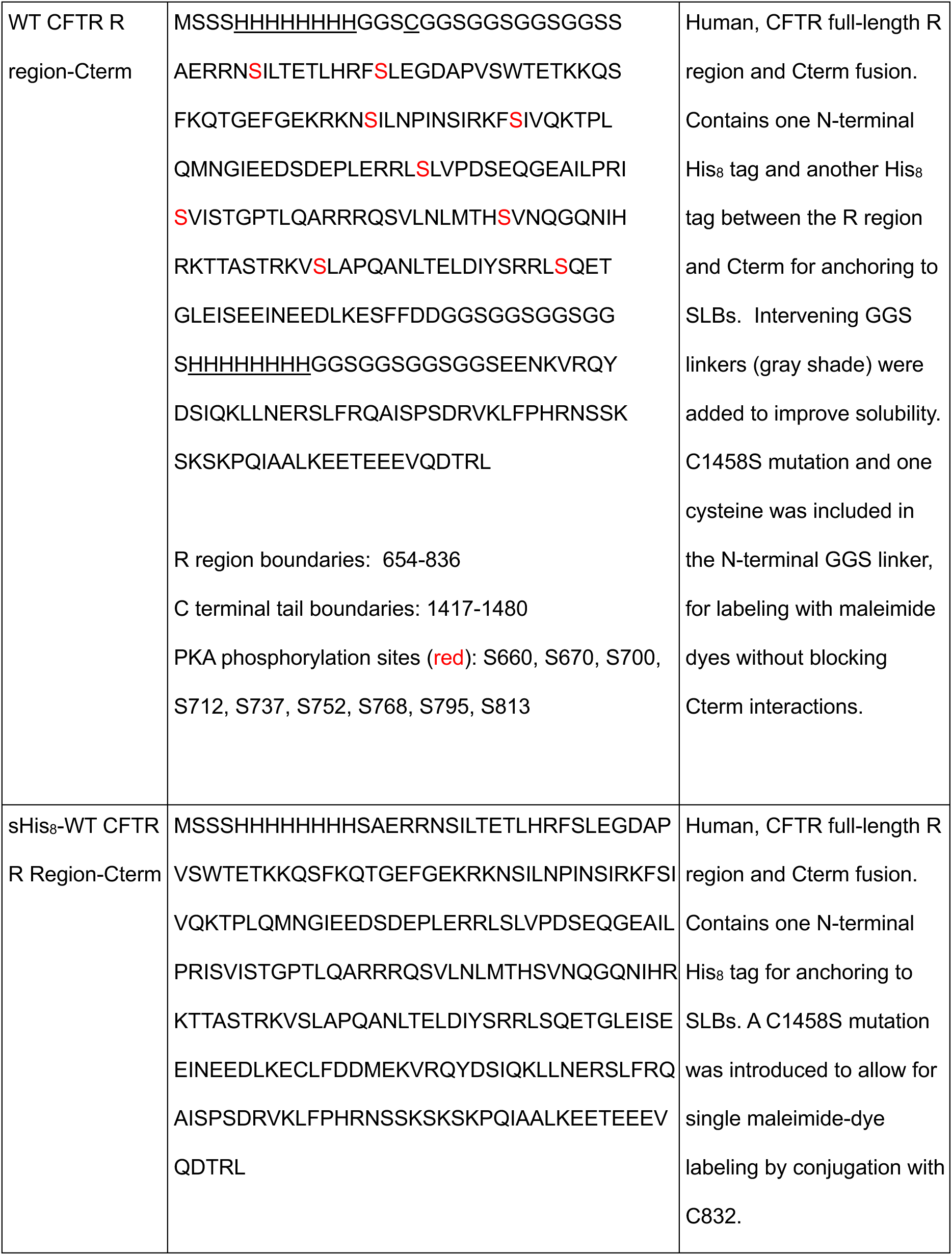
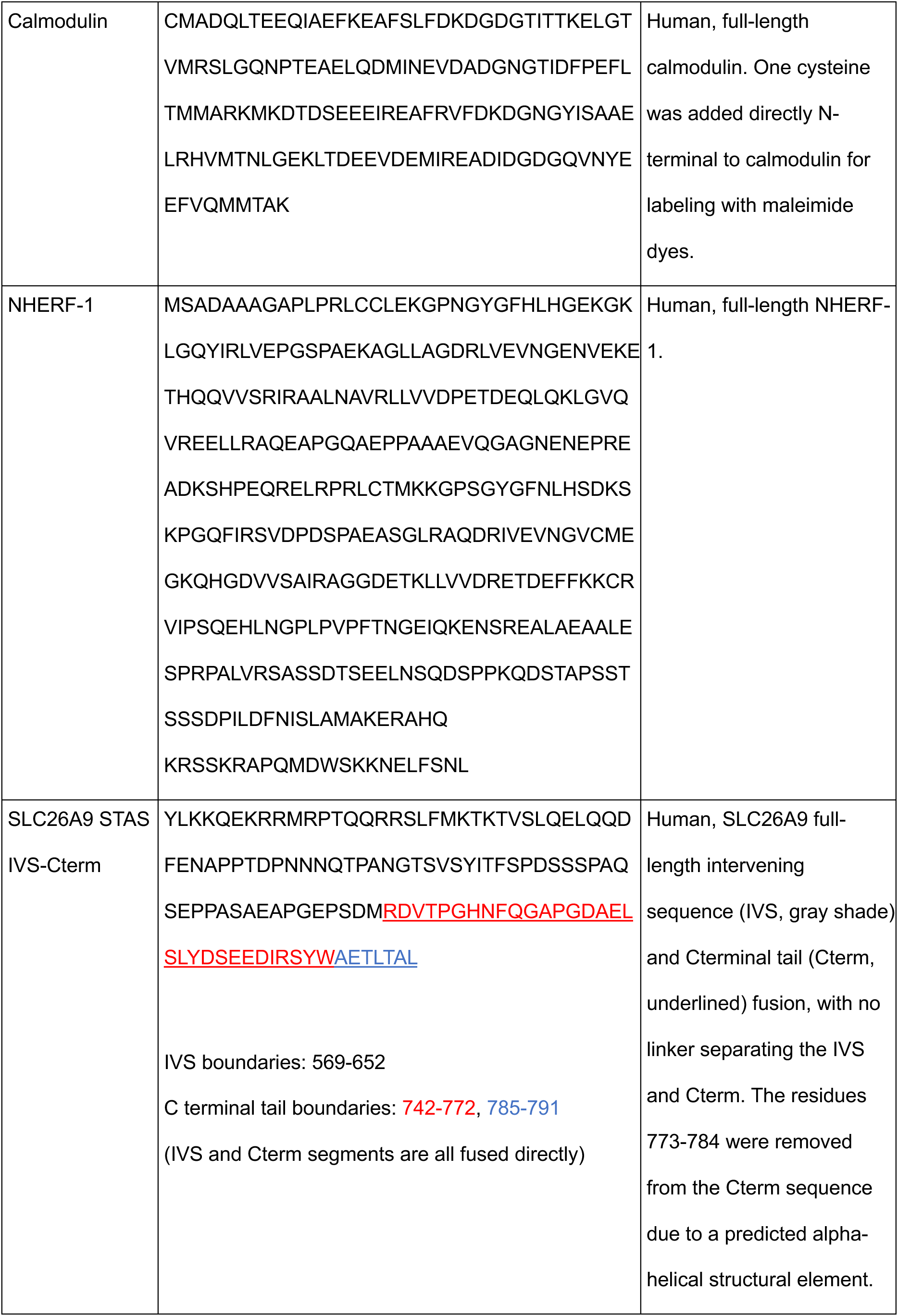
Amino acid sequences of all recombinant proteins used in this study.

### Protein expression, purification, and maleimide dye labeling

All proteins were expressed in BL21-CodonPlus (DE3)-RIPL Competent *E. coli* bacteria transformed with plasmids encoding the genes described previously. Bacteria were grown in lysogeny broth (LB) containing 50 mg/mL kanamycin and 34 mg / ml chloramphenicol until O.D._600_ = 0.6-0.8. Protein expression was induced using 1 mM isopropyl β-D-1-thiogalactopyranoside (IPTG) and bacteria were incubated at 16 °C for 16-18 hours before harvesting by centrifugation at 4500 rpm for 30 minutes at 4 °C. Cell pellets were either used immediately or frozen and stored at −80 °C for up to 2 months before purification. Purification protocols were adapted from previously published methods [31,32] and are detailed below. All proteins were verified by mass spectrometry analysis.

#### Purification of CFTR R region-Cterm fusions and SLC26A9 STAS IVS-Cterm

The CFTR R region-Cterm and His_6_-SUMO-SLC26A9 STAS IVS-Cterm constructs were purified from the insoluble lysate fraction. Pellets were resuspended in 20 ml guanidinium chloride buffer (6M GdmCl, 50 mM Tris-HCl (pH 7.5), 150 mM NaCl, and 0.2 mM TCEP) per liter of culture. Cell resuspensions were lysed by sonication on ice for 10 minutes (30% amplitude, cycled at 2 secs on / 2 secs off) and clarified by centrifugation at 8500 RPM for 1 hour using a fixed angle JA-20 rotor (Beckman Coulter) at 4 °C. Supernatant fractions were loaded onto HisPur Ni-NTA resin (Thermo Scientific) as per manufacturer’s instructions and washed with 5 CVs of GdmCl buffer containing 20mM imidazole (pH 7.5), and eluted in 300 mM imidazole (pH 7.5) in 5 x1 CV fractions. Aliquots of fractions containing protein were precipitated from GdmCl by trichloroacetic acid precipitation and resuspended in 8M urea before analysis by gel electrophoresis. Proteins were further purified by size exclusion chromatography using a HiLoad 26 / 600 Superdex 75 prepgrade column (Cytiva), in GdmCl buffer. SLC26A9 STAS IVS-Cterm was diluted to 240 mM GdmCl, incubated with 10 units of ULP-1 per µg of protein for 4 hours at room temperature, and flowed over Ni-NTA to remove the His_6_-SUMO tag. Gels confirming fusion protein purifications are shown in Supplemental Figure S6B.

#### Purification of calmodulin and NHERF-1

His_6_-SUMO-Calmodulin and His_6_-SUMO-NHERF-1 were purified from the soluble lysate fraction following the previously described method for CFTR R region-Cterm and SLC26A9 STAS IVS-Cterm constructs, except with non-denaturing Tris buffer (20 mM Tris-HCl (pH 7.5), 150mM NaCl, and 0.2 mM TCEP). Protein-loaded Ni-NTA resin was washed stepwise using Tris buffers containing 20mM imidazole (pH 7.5), and eluted in 300mM imidazole (pH 7.5). Calmodulin and NHERF-1 were incubated in Tris buffer with 10 units of ULP-1 per 20 µg of protein for 4 hours at room temperature and flowed over Ni-NTA to remove the His_6_-SUMO tag. Gels confirming fusion protein purifications are shown in Supplemental Figure S6B.

#### Labeling CFTR R region-Cterm with maleimide dye

This labeling protocol was adapted from previously published methods [59,72]. βME was added to a final concentration of 10 mM to purified CFTR R region-Cterm to reduce all cysteine residues. CFTR R region-Cterm was then buffer exchanged into a buffer of 3M GdmCl, 50 mM HEPES-NaOH (pH 7.0), and 150 mM NaCl, and incubated with 3-times molar excess of Alexa Fluor (AF) 647 C2-maleimide conjugated dye (Invitrogen) for 16 hours at 4 °C in the dark. The labeling reaction was quenched with 10 mM βME, and excess dye was removed by size exclusion chromatography using a 10 / 300 Superdex 75 GL column (Cytiva).

#### Phosphorylation of Model CFTR R region-Cterm

This phosphorylation protocol was adapted from a previously published method [32]. Purified and AF647-labelled CFTR R region-Cterm was diluted to below 100 mM GdmCl. 100 units of protein kinase A (PKA) was added per 500 µg of CFTR R region-Cterm and dialyzed in a buffer of 50 mM Tris-HCl (pH 7.5), 150 mM NaCl, 20 mM MgCl_2_, 10 mM ATP, and 2 mM DTT for 16 hours at 4 °C in the dark. The buffer was exchanged for fresh buffer midway through the reaction to refresh the ATP. The phosphorylation reaction was flowed over Ni-NTA to remove His-tagged PKA and confirmed by mass spec analysis.

### Biolayer interferometry

All measurements were performed on a FortéBio Octet RED384 instrument using Octet Ni-NTA biosensors (Sartorius) and 384-well black microplates with either flat (Greiner) or tilted bottoms (Sartorius). Biosensors were hydrated in Sartorius Kinetics Buffer for 5 minutes prior to use. All protein was initially dialyzed into HEPES buffer (50 mM HEPES (pH 7.3), 150 mM NaCl, and 1 mM TCEP) and freshly diluted in Kinetics Buffer prior to use. All biolayer interferometry experiments were performed as follows: biosensors were incubated in Kinetics Buffer for 120 seconds (Baseline), 500 nM CFTR R region-Cterm for 120 seconds (Loading), BSA Blocking buffer for 60 seconds (Quenching), analyte (CFTR R region-Cterm, Calmodulin, NHERF-1, or SLC26A9 STAS IVS-Cterm) for 300-600 seconds (Association), and Kinetics Buffer for 300 seconds (Dissociation). Biosensors were regenerated up to 10 times by alternately washing in Kinetics Buffer containing 750 mM and 0 mM imidazole (pH 7.5). To minimize evaporation effects, a working volume of 75 µL per well was used and experimental runs did not exceed 1 hour. All measurements were performed in triplicate. Binding curves were processed and fit using Octet Analysis Studio Software (Sartorius).

### Reconstitution assays using supported lipid bilayers

The methods described in this section were adapted from previously published methods [17,57,59,71,72].

#### Preparation of small unilamellar vesicles (SUVs)

Synthetic phospholipids and cholesterol were all sourced from Avanti Polar Lipids, barring 16:0 DGS-(Ni)NTA, which was synthesized and provided by the Levental lab. Lipids for reconstitution experiments were combined as detailed in Table 3, dried under a stream of Argon, desiccated for 12 hours, and resuspended in phosphate-buffered saline (pH 7.3) to create solutions of multilamellar lipid vesicles. Lipid solutions were repeatedly frozen in liquid nitrogen and thawed on a 45 °C heat block 30-45 times, or until the solution cleared, to generate small unilamellar vesicles (SUVs). Lipid solutions were centrifuged at 21,000 g for 2 hours at 4 °C to pellet any remaining multilamellar vesicles. Supernatant containing SUVs was collected, topped with Argon, and stored at 4 °C for up to 2 weeks. Lipid solutions were frozen and stored after 15 freeze-thaws at −80 °C for up to 6 months.

#### Preparation of supported lipid bilayers (SLBs)

Supported lipid bilayers (SLBs) were formed on 96-well black glass-bottom plates (Corning). Glass was washed with 5% Hellmanex III at 50 °C for 3 hours, thoroughly rinsed with MilliQ H_2_O, washed twice with 6M NaOH at 45°C for 1 hour, and thoroughly rinsed with MilliQ H_2_O, before equilibrating in HEPES buffer (50 mM HEPES (pH 7.3), 150 mM NaCl, and 1 mM TCEP). SUVs were added to wells containing HEPES buffer and incubated at 40 °C for 45 min to allow SUVs to collapse and fuse on the glass substrate to form SLBs. SLBs were washed with HEPES buffer three times to remove excess SUVs. For double SLBs, base layer SUVs were incubated to form the base SLB, washed three times with HEPES buffer, then incubated with SUVs composed of lipids for the top bilayer for 1 hour at 40 °C, followed by a second wash step with HEPES buffer, as we have previously described [72]. Table 3 contains information describing the composition of supported lipid bilayers.

HEPES buffer containing 1 µM CFTR R region-Cterm protein was incubated on the prepared SLBs for 1 hour at room temperature, resulting in a final CFTR R region-Cterm density of approximately 5000 molecules / µm^2^ on SLBs and (d)SLBs. (d)SLBs were washed three times with HEPES buffer. For conditions containing CFTR interactors, proteins were incubated on (d)SLBs coated with CFTR R region-Cterm proteins for 15-30 minutes at the following concentrations: 1 µM calmodulin, 0.5 µM NHERF-1, and 1 µM SLC26A9 STAS IVS-Cterm. For calcium-containing conditions, 0.5 mM CaCl_2_ was added to the HEPES buffer, and interactors were dialyzed into HEPES buffer containing the same concentration of calcium. All experiments were performed in triplicate.

#### Evaluation of membrane fluidity with fluorescence recovery after photobleaching

Fluorescence recovery after photobleaching (FRAP) assays were used to verify the fluidity of (d)SLBs and the proteins anchored to them. All FRAP experiments were performed on a point scanning Leica SP8 LIGHTNING Confocal Microscope (Leica Microsystems) equipped with Leica Application Suite X software, using a 63x / 1.3 NA oil immersion objective. 488 and 638 nm lasers were used at 100 % power for 0.65 seconds to bleach AF488-labelled (d)SLBs (18:1 PE-TopFluor AF488, Avanti cat. #810386) or AF647-labelled proteins, respectively. (d)SLBs and proteins were bleached at a small region of interest (ROI) to 40 % fluorescence intensity or lower at room temperature, and recovery within the ROI was monitored for 150 – 300 seconds. Following photobleaching, changes in fluorescence intensity of the ROI over time produce the recovery curve. Due to gradual photobleaching of the entire microscope field of view during imaging, FRAP recovery curves were corrected by normalizing ROI fluorescence intensity to background fluorescence intensity over time.

#### Imaging and analysis of CFTR R region-Cterm clusters

Bilayer images were obtained on a point scanning Leica SP8 LIGHTNING Confocal Microscope (Leica Microsystems) equipped with Leica Application Suite X software. Images were obtained using a 63x / 1.3 NA oil immersion objective with 488 and 638 nm lasers.

Image analysis was performed using Fiji. All images were processed at identical brightness and intensity settings. The degree of clustering was determined by measuring the variance of intensities (Standard Deviation^2^) within each image, as we previously described [71]. T-test analyses were performed, and data were plotted using GraphPad Prism (Dotmatics).

## ACKNOWLEDGEMENTS

We thank the Levental Lab at the University of Virginia for the generous gift of 16:0 DGS-NTA(Ni) used to target His-tagged proteins to cholesterol-rich membrane environments. We thank Drs. Andrew Chong, Christine Bear, Trevor Moraes, and Hyun Lee for stimulating discussions, the Hospital for Sick Children Structural & Biophysical Core Facility and the Hospital for Sick Children Imaging Facility for use of the instruments used in this study, and Drs. Leslie Loew and Michael Blinov for their assistance using the Virtual Cell which is supported by NIH Grant R24 GM137787 from the National Institute for General Medical Sciences. YW was supported by a Canada Graduate Scholarship – Master’s Award from the Natural Sciences and Engineering Research Council. JDF-K acknowledges support from Cystic Fibrosis Canada (#604926) and the Canada Research Chairs Program. JAD acknowledges support for studies of membrane-associated phase separation from Natural Sciences and Engineering Research Council Discovery Grant (RGPIN-2022-03274);

## AUTHOR CONTRIBUTIONS

YW, JDF-K, and JAD designed the study. YW, RH, and JS prepared reagents and performed experiments. YW analyzed data. YW, JDF-K, and JAD wrote and edited the manuscript with comments from RH and JS.

## SUPPLEMENTAL FIGURE LEGENDS

**Figure S1. A)** Visualized map of modeled interaction network using the Virtual Cell. “Reacting” and “Reactions” here refer to binding interactions.

**Figure S2.** Modeled interactions between domains and binding motifs in CFTR and its binding partners. Green lines indicate protein-protein interaction affinities defined by our study using BLI. Magenta lines indicate protein-protein, protein-cholesterol, and cholesterol-cholesterol affinities published in the literature. R1, R2, R3, and R4 indicate distinct CFTR R regions that can interact with binding partners, as determined by our previous interaction analysis using NMR [31,32]. For each binding motif, we used the overall affinity measured by BLI, as we have previously done for Nck: N-WASP interactions [58,91].

**Figure S3. A)** Images of AlexaFluor (AF) 488-labelled sHis_8_-CFTR R region-Cterm on supported lipid bilayers (SLBs) in 0 mM (left) or 275 mM (right) imidazole-containing buffer. Scale bar = 20 µm. **B)** SLBs composed of 15% cholesterol fully recover following photobleach (top). SLBs composed of 20% cholesterol do not recover (bottom). n = 5. Scale bar = X µm. **C)** The AlexaFluor488 fluorescence of the bottom SLB of (d)SLBs recovers after photobleaching to the same extent as an uncovered SLB indicating that layering a second SLB on top of the SLB contacting the coverglass subtrates has no effect on fluidity (left). n = 5. The fluorescence of the upper SLB of (d)SLBs recovers fully from when composed of 20 – 40% cholesterol indicating that it remains fully fluid, unlike its SLB counterpart **(B)**. n = 5.

**Figure S4. A)** Sequences of double-His_8_-tagged WT and Model CFTR R region-Cterm constructs. NMR-detected regions of protein-protein interaction were fused with GGS linkers in Model CFTR R region-Cterm. This allows us to parse the impact of specific protein - protein and protein – lipid interactions. Dual His-tags were used to anchor the fusion protein to supported lipid bilayers in lieu of transmembrane domains. **B)** Schematics illustrating NMR-determined binding elements within the CFTR R region from [31,32]. These elements were included in our Model CFTR R region-Cterm fusion protein shown in **(A)** and Figure S6, while all residues residing outside these interaction elements were replaced with GGS linkers. (Row 1) Schematic of sequence for wild type CFTR R region shown in **(A)**. (Row 2) Schematic of sequence for Model CFTR R region shown in **(A).** (Row 3) Schematic showing calmodulin-binding elements in the presence of Ca^2+^ between R region residues 659-671, 700-715, and 760-780 in the non-phosphorylated R region. (Row 4) Schematic showing NMR-derived binding elements of the R region to the SLC26A3 STAS domain including its IVS between residues 660-681, 750-777, 805-814, and 829-836 with and without phosphorylation of the R region. In the previously published NMR experiments [31], the homologous SLC26A3 STAS domain with IVS was used due to instability of the STAS with IVS of SLC26A9. As shown in Figure S6, here we use a fusion of only the IVS (lacking the folded STAS domain) together with the Cterm of SLC26A9 for our biochemical assays. (Row 5) Schematic showing Cterm binding elements between residues 663-680, 708-723, 750-778, and 803-815 in the non-phosphorylated R region. (Row 6) Schematic showing Cterm binding elements between residues 669-695, 708-723, 750-778, 802-810, and 827-836 in the phosphorylated R region.

**Figure S5.** Schematic of double supported lipid bilayers ((d)SLB) used in experiments shown in Figures 5, 6, and 7. **A)** (d)SLB composed of cholesterol-rich and -depleted domains with CFTR-R region-Cterm anchored to 18:1 DGS-NTA(Ni) in cholesterol-depleted domains. **B)** (d)SLBs composed of cholesterol-rich and -depleted domains with CFTR-R region-Cterm anchored to 16:0 DGS-NTA(Ni) in cholesterol-rich domains.

**Figure S6.** Schematics, including residue positions, **(A)** and Coomassie stained gels **(B)** of recombinant fusion proteins used in this study.

