## Supplementary figures and images for "Protein interactions, calcium, phosphorylation, and cholesterol modulate CFTR cluster formation on membranes"

### Supplemental Material

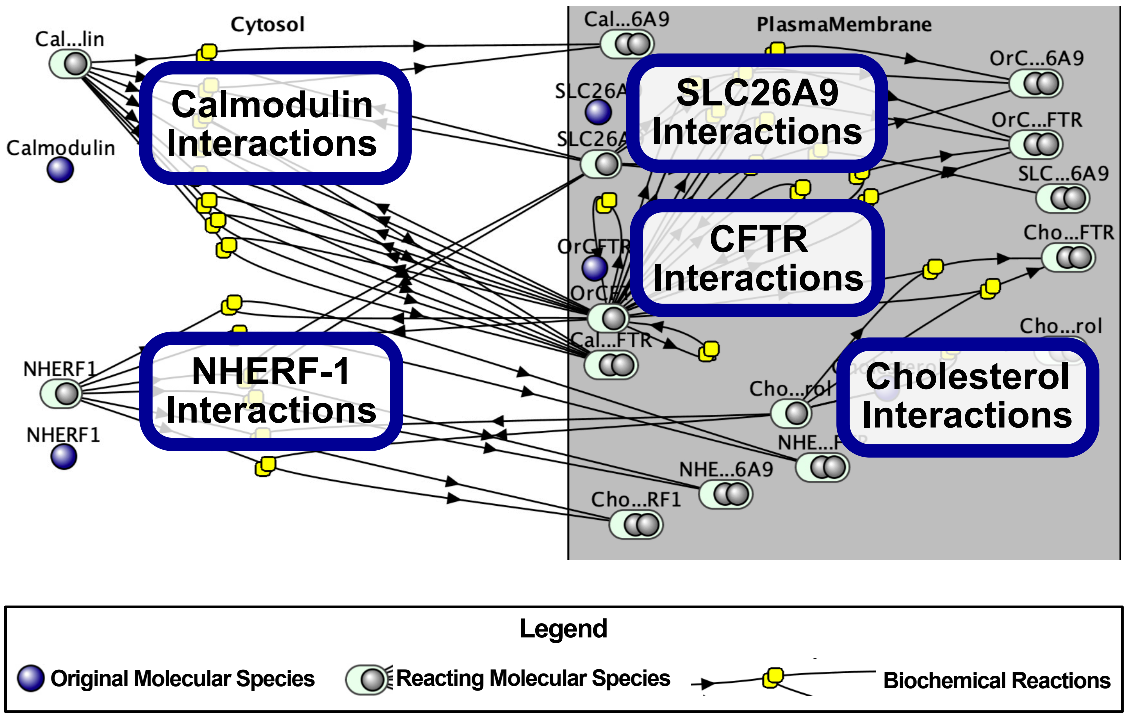

#### Molecular interactions modeled using the Virtual Cell

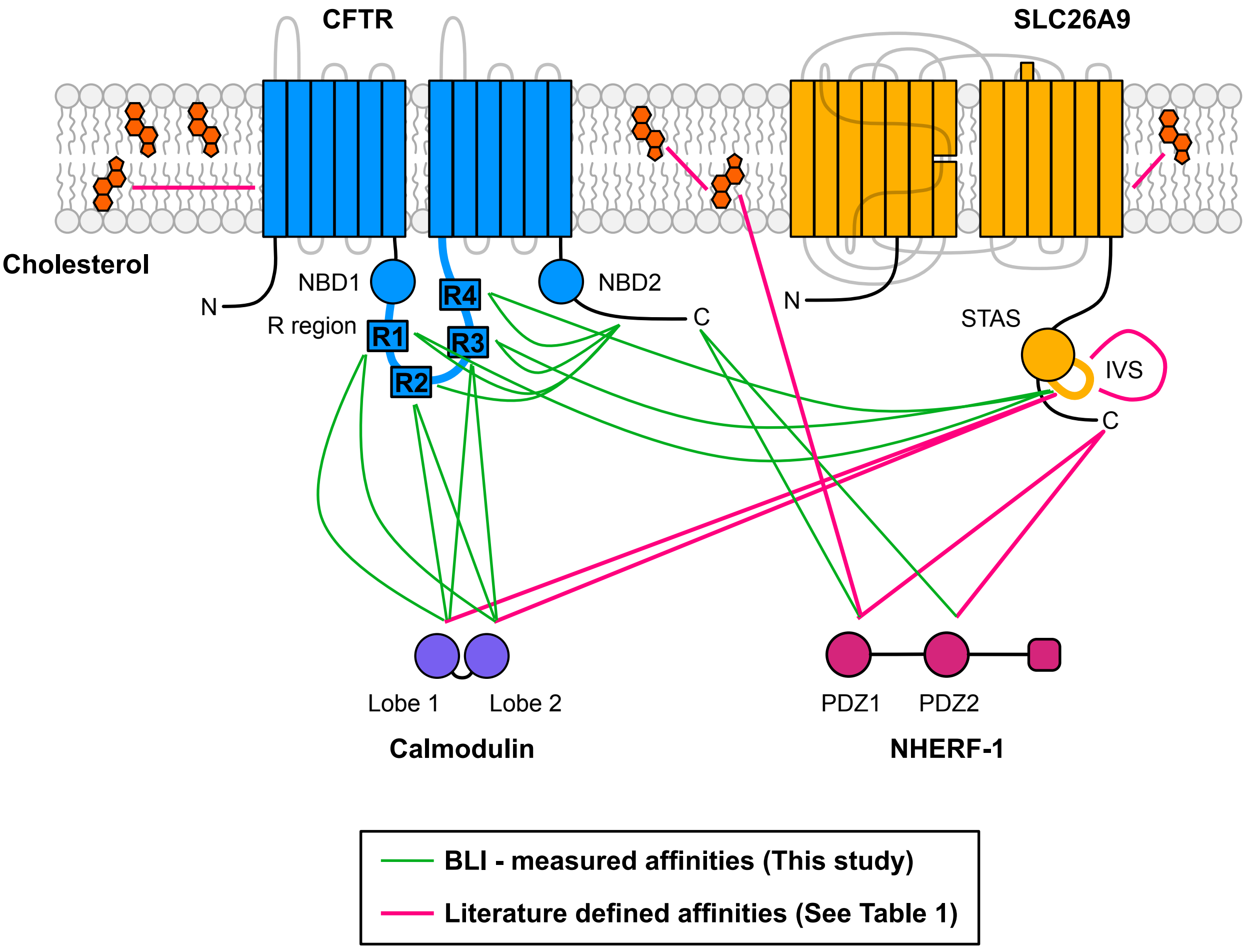


Figure S2

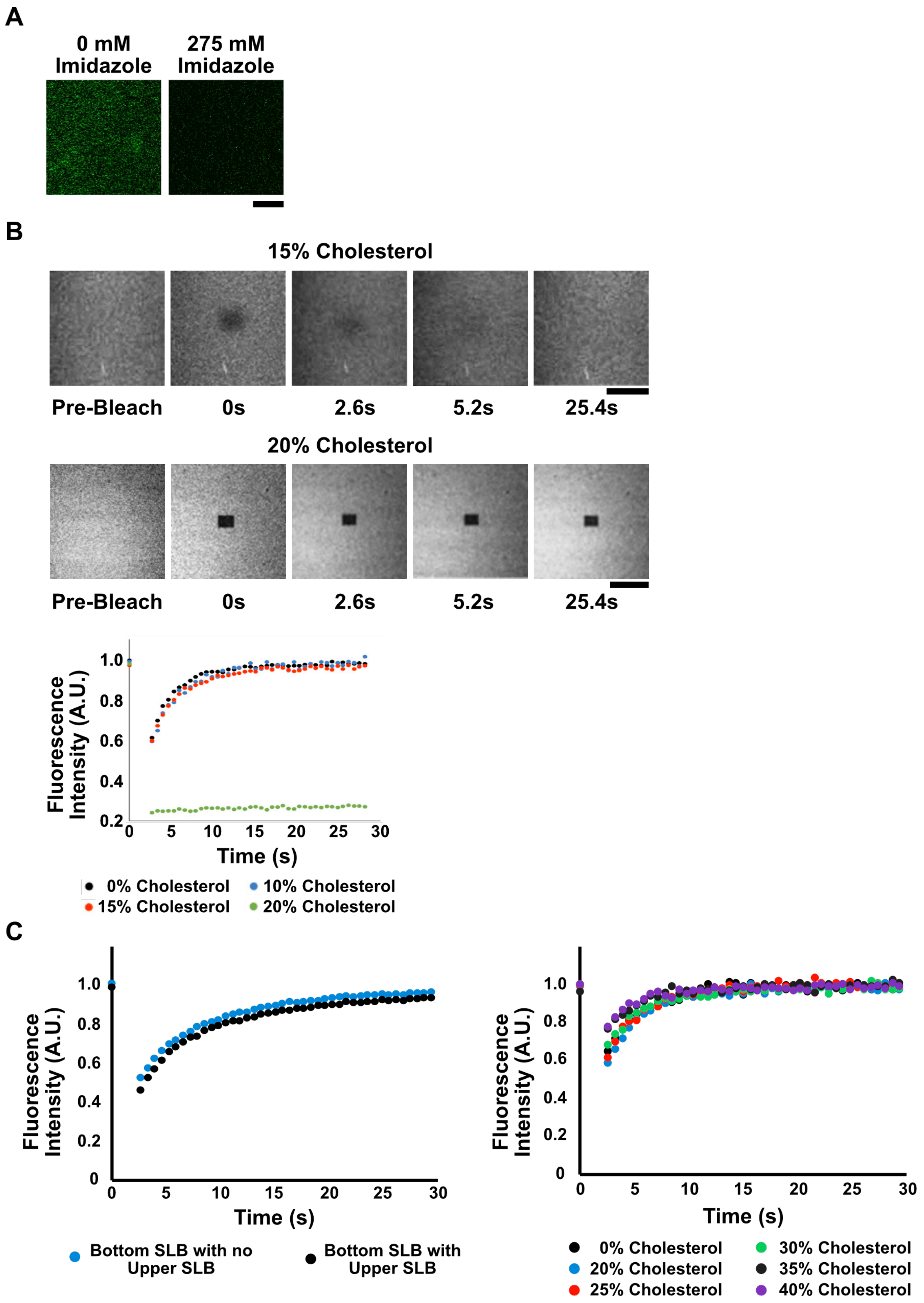

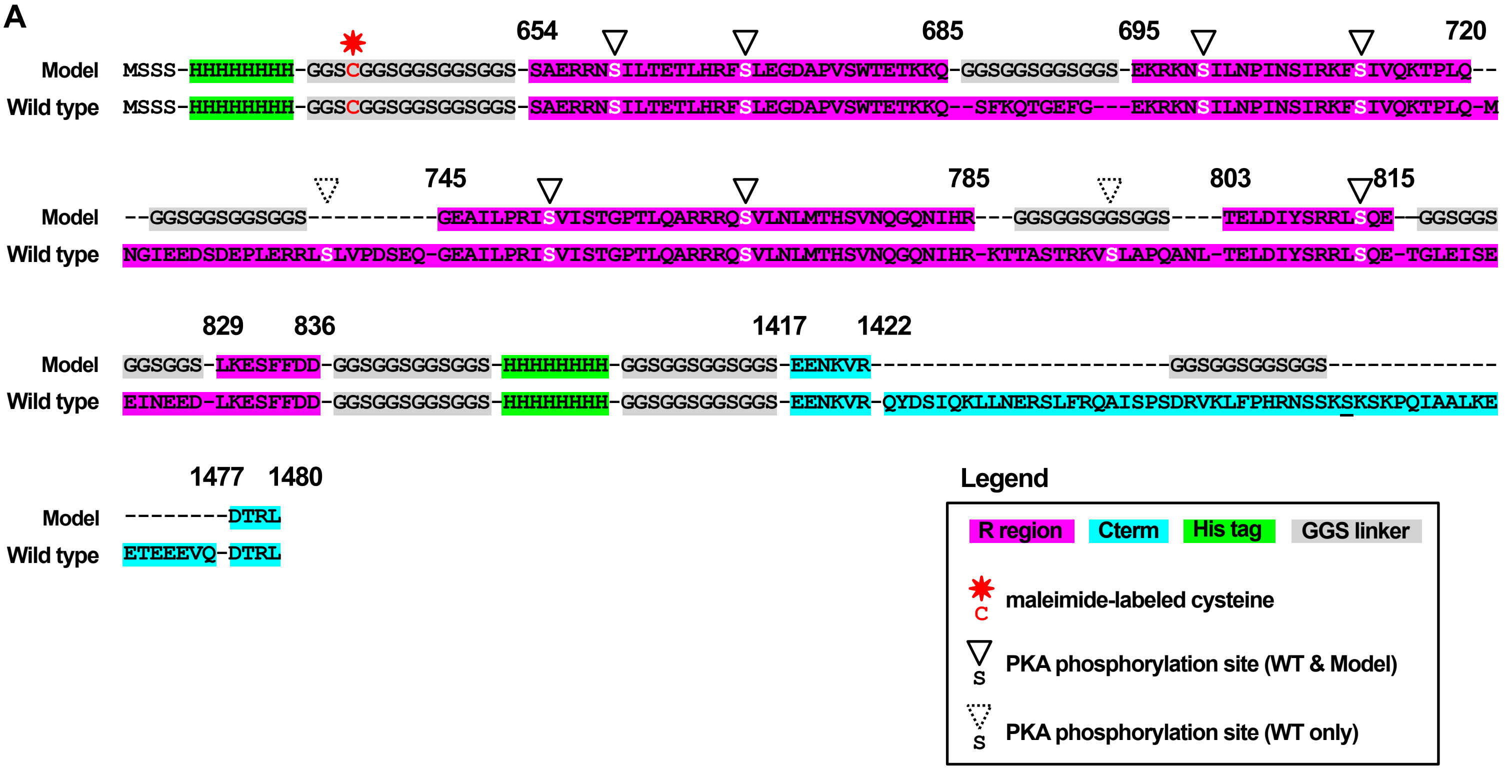

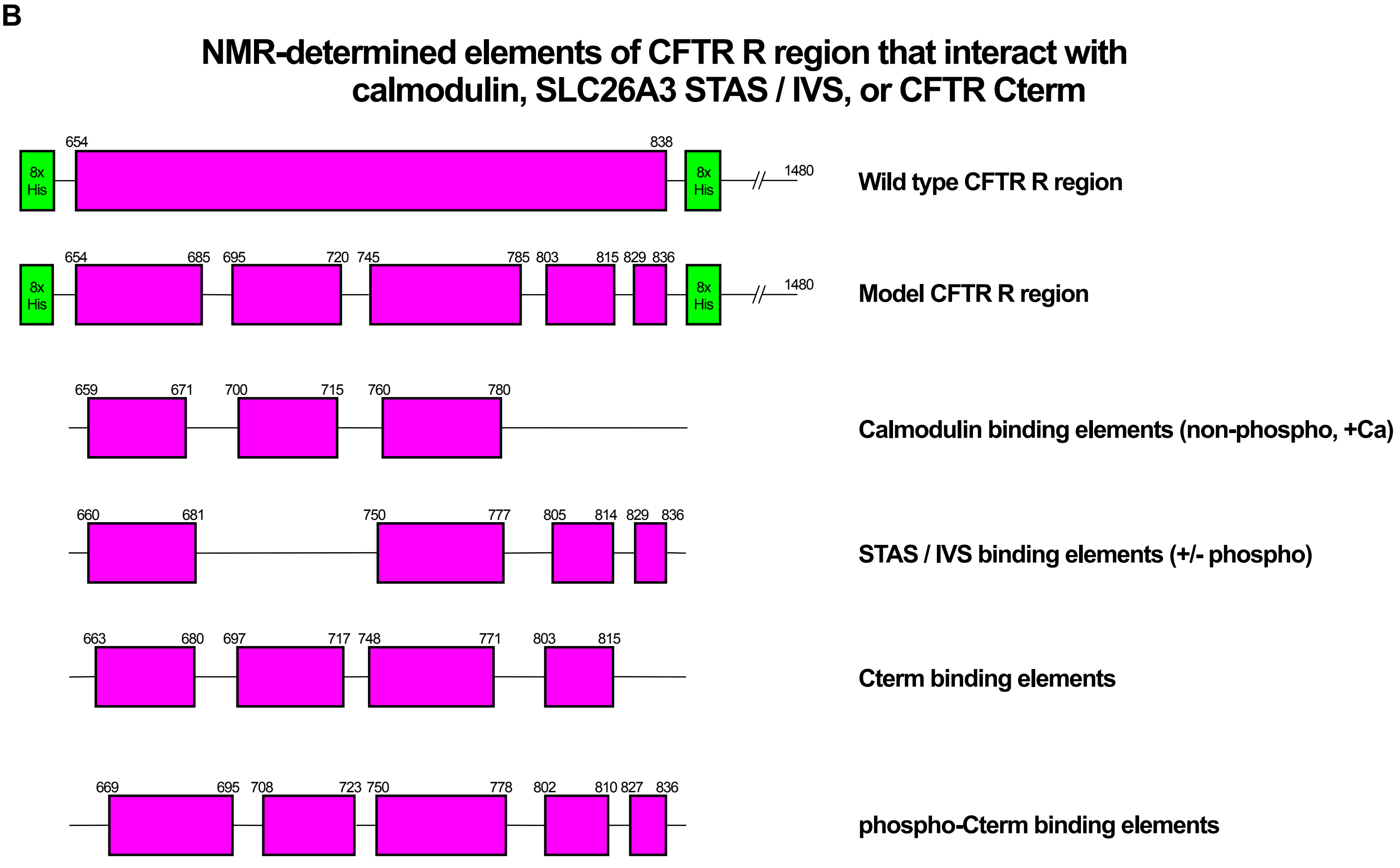

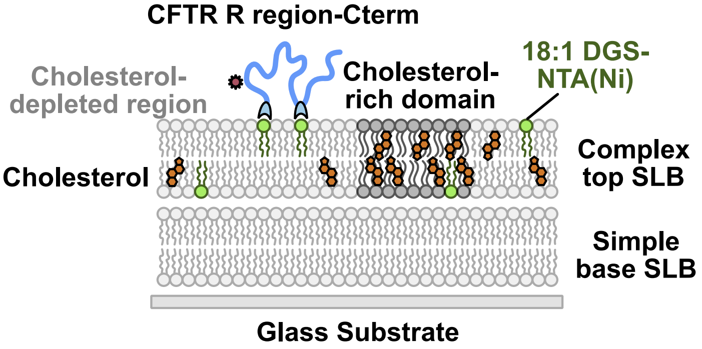

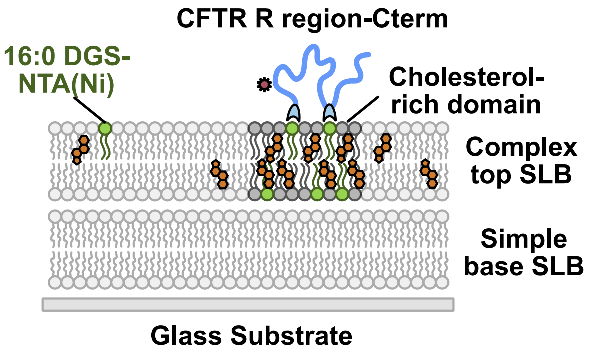

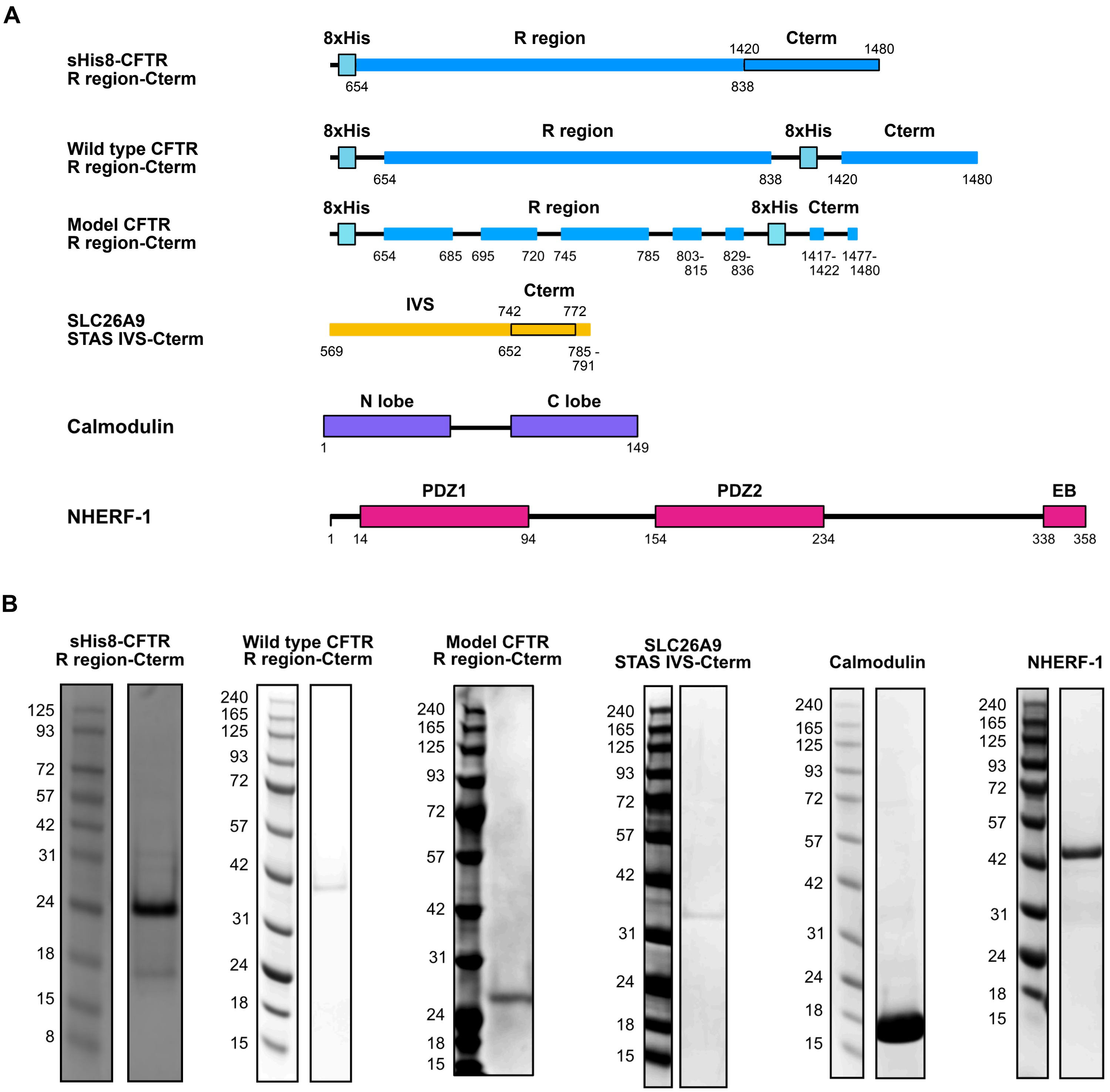
